# Systematic mapping of chromatin state landscapes during mouse development

**DOI:** 10.1101/166652

**Authors:** David U. Gorkin, Iros Barozzi, Yanxiao Zhang, Ah Young Lee, Bin Li, Yuan Zhao, Andre Wildberg, Bo Ding, Bo Zhang, Mengchi Wang, J. Seth Strattan, Jean M. Davidson, Yunjiang Qiu, Veena Afzal, Jennifer A. Akiyama, Ingrid Plajzer-Frick, Catherine S. Pickle, Momoe Kato, Tyler H. Garvin, Quan T. Pham, Anne N. Harrington, Brandon J. Mannion, Elizabeth A. Lee, Yoko Fukuda-Yuzawa, Yupeng He, Sebastian Preissl, Sora Chee, Brian A. Williams, Diane Trout, Henry Amrhein, Hongbo Yang, J. Michael Cherry, Yin Shen, Joseph R. Ecker, Wei Wang, Diane E. Dickel, Axel Visel, Len A. Pennacchio, Bing Ren

**Affiliations:** Ludwig Institute for Cancer Research, 9500 Gilman Drive, La Jolla, CA 92093-0653, USA; University of California, San Diego School of Medicine, Department of Cellular and Molecular Medicine, Institute of Genomic Medicine, and Moores Cancer Center, 9500 Gilman Drive, La Jolla, CA 92093-0653, USA; Functional Genomics Department, Lawrence Berkeley National Laboratory, 1 Cyclotron Road, Berkeley, California 94720, USA; U.S. Department of Energy Joint Genome Institute, Walnut Creek, California 94598, USA.; School of Natural Sciences, University of California, Merced, Merced, California 95343, USA.; Dept. of Biochemistry and Molecular Biology, Penn State School of Medicine, 500 University Drive, MC H171, Hershey, PA 17033, USA.; Institute for Human Genetics and Department of Neurology, University of California, San Francisco, San Francisco, California 94143, USA; Bioinformatics and Systems Biology Graduate Program, University of California, San Diego, La Jolla, California 92093; Genomic Analysis Laboratory and Howard Hughes Medical Institute, Salk Institute for Biological Studies, La Jolla, California 92037; Howard Hughes Medical Institute, Salk Institute for Biological Studies, La Jolla, California 92037; Stanford University School of Medicine, Department of Genetics, Stanford, California, USA; Division of Biology and Biological Engineering, California Institute of Technology, Pasadena, CA 91125, USA; Comparative Biochemistry Program, University of California, Berkeley, CA 94720, USA

**Author notes:** equal contribution (David U. Gorkin, Iros Barozzi, Yanxiao Zhang).

## Abstract

Embryogenesis requires epigenetic information that allows each cell to respond appropriately to developmental cues. Histone modifications are core components of a cell’s epigenome, giving rise to chromatin states that modulate genome function. Here, we systematically profile histone modifications in a diverse panel of mouse tissues at 8 developmental stages from 10.5 days post conception until birth, performing a total of 1,128 ChIP-seq assays across 72 distinct tissue-stages. We combine these histone modification profiles into a unified set of chromatin state annotations, and track their activity across developmental time and space. Through integrative analysis we identify dynamic enhancers, reveal key transcriptional regulators, and characterize the role of chromatin-based repression in developmental gene regulation. We also leverage these data to link enhancers to putative target genes, revealing connections between coding and non-coding sequence variation in disease etiology. Our study provides a compendium of resources for biomedical researchers, and achieves the most comprehensive view of embryonic chromatin states to date.

## MAIN TEXT

Embryonic development relies on a complex interplay between genetic and epigenetic factors. While genetic information encoded in DNA sequence provides the instructions for an embryo to develop, epigenetic information is required for each cell type in an embryo to obtain its specialized function from a single set of instructions. More specifically, the functional output of a given DNA sequence depends on its chromatin state, which is defined in large part by posttranslational modifications of histone proteins^1,2^. These histone modifications take many forms, and they can influence gene expression by directly modulating DNA-histone interactions, or serving as binding sites for cognate reader proteins that function downstream to enhance or repress transcription^3^. Importantly, histone modifications allow for the transmission of gene activity states through cell division^4^, which is crucial for the maintenance of cellular identity. Given the importance of histone modifications in gene regulation, it is not surprising that mutations in genes that encode histones or histone modifying enzymes can lead to a wide variety of diseases including cancer, neurodevelopmental disorders, and congenital malformations^5-7^.

Histone modifications have also proven to be valuable markers for genome annotation^8,9^. Different types of functional sequences have distinct patterns of histone modifications, which can be profiled using Chromatin Immunoprecipitation sequencing (ChIP-seq). For example, chromatin around active promoters is characterized by H3K4me3 and H3K27ac, while chromatin around active enhancers is often marked by H3K4me1 and H3K27ac^10-12^. While the profiles of individual histone modifications can be informative, the most comprehensive functional annotations are achieved when multiple histone modifications are integrated into a unified set of chromatin states based on combinatorial patterns^13-15^. Each region in the genome can then be classified according to its chromatin state, providing insight into its potential function.

### Profiling of histone modifications across developmental time and tissues

Despite the importance of chromatin states in determining the functional output of the genome, a comprehensive survey of chromatin states during mammalian embryonic development has been lacking. The mouse is an ideal system to study the developmental epigenome because it is the most widely used animal model in biomedical research, and offers experimental access to early stages of embryonic development. We harvested mouse tissues at closely spaced intervals from 10.5 days post conception (E10.5) until birth, which is roughly equivalent to gestational week 3 of human development through the end of the first trimester of pregnancy. At each stage, we dissected a diverse panel of tissues from multiple litters of C57BL/6N embryos and performed 2 replicates of ChIP-seq for each of 8 histone modifications (H3K4me1, H3K4me2, H3Kme3, H3K27ac, H3K27me3, H3K9ac, H3K9me3, H3K36me3) (Figure 1a-b; Extended data figure 1). These include all six “core” histone modifications required by the International Human Epigenome Consortium (IHEC) to generate a reference epigenome^16,17^. At the earliest stage sampled (E10.5) it was not feasible to harvest enough tissue for standard ChIP-seq, so we used a modified micro-ChIP-seq procedure designed to work on much smaller numbers of cells and restricted our scope to only these 6 histone modifications. The complete data series includes more than 66 billion sequencing reads from 564 ChIP-seq experiments, each consisting of two biological replicates derived from different embryo pools (N=1,128 replicates total). All datasets were generated according to ENCODE standards and processed with a standardized pipeline that only outputs peaks passing strict reproducibility criteria^18^ (see **Methods**; Extended data figure 2a). In total, we identified more than 45 million peaks covering roughly one-third of the mouse genome (Extended data figure 2b-d).

**Figure 1:**
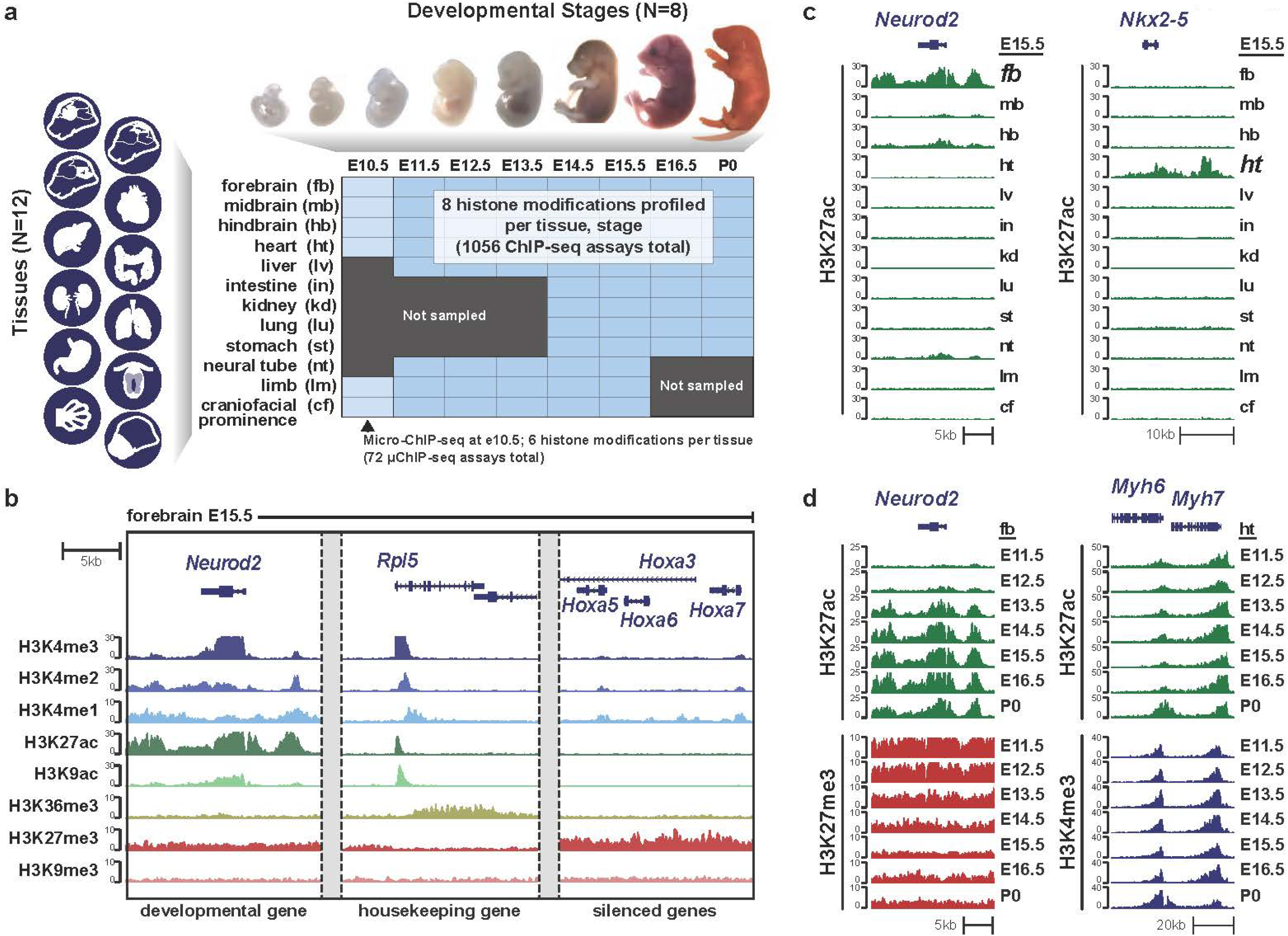
Profiling histone modifications during mouse embryonic development. (a) Overview of the experimental design. (b) Characteristic patterns of histone modifications at different types of gene loci: *Neurod2* (chr11:98,318,134-98,336,928; mm10), *Rpl5* (chr5:107,895,370-107,914,164; mm10), *Hoxa* (chr6:52,199,870-52,218,665; mm10). Data shown are from forebrain at E15.5. (c) Tissue-restricted H3K27ac enrichment at tissue-restricted regulators *Neurod2* and *Nkx2-5* (chr17:26,832,795-26,855,694; mm10). Data shown from 12 tissues at stage E15.5. (d) Stage-restricted histone modification patterns at developmentally regulated loci *Neurod2* and *Myh6*/*Myh7* (chr14:54,927,121-55,010,762; mm10). Note the switch in activity from *Myh7* to *Myh6* at P0.

We observed several high-level features of the data series that warrant comment here. First, as expected, the landscape of histone modifications is highly variable between tissues, particularly for marks of activity such H3K27ac and H3K9ac (Figures 1c, 2a; Extended data figure 3d). We observe that tissues with closer developmental origins tend to have similar histone modification profiles. For example, tissues of the Central Nervous System (CNS) can be easily distinguished from other tissues by hierarchical or k-means clustering (Figures 2a-b), as well as by principle component analysis (Extended data figure 2e). In addition to differences between tissues, histone modification profiles change progressively within each tissue during development (Figures 1d, 2c-d, Extended data figure 3). These developmental dynamics likely reflect at least two underlying biological processes: 1) changes in the epigenetic landscape of individual cells within a tissue as they undergo differentiation, and 2) shifts in the relative abundance of different cell types that compose a tissue. Although we cannot separate the contributions of these two factors, many of the epigenetic changes we observe reflect known hallmarks of cellular differentiation. For example, we observe a dramatic shift in activity from *Myh7* to *Myh6* in heart at P0, which is known to occur in cardiomyocytes just prior to birth^19^ (Figure 1d). In the developing forebrain, we observe that markers of mature neurons such as *NeuroD2* and *Gad1* acquire active histone modifications during development, and concomitantly lose repressive modifications^20,21^ (Figure 1d; Extended data figure 3a). We observe the opposite trend at genes encoding key cell cycle factors such as *Ccnb1* and *Cdk2*, where active histone modifications decrease during forebrain development as neurons mature and exit the cell cycle (Extended data figure 3b-c). As a general trend, we observe that regions of histone modification restricted in developmental time or space are more likely to be distal to Transcriptional Start Sites (TSS; Extended data figure 3d). This is consistent with the model that TSS-distal regulatory elements play a key role in determining cellular identity^22,23^.

**Figure 2:**
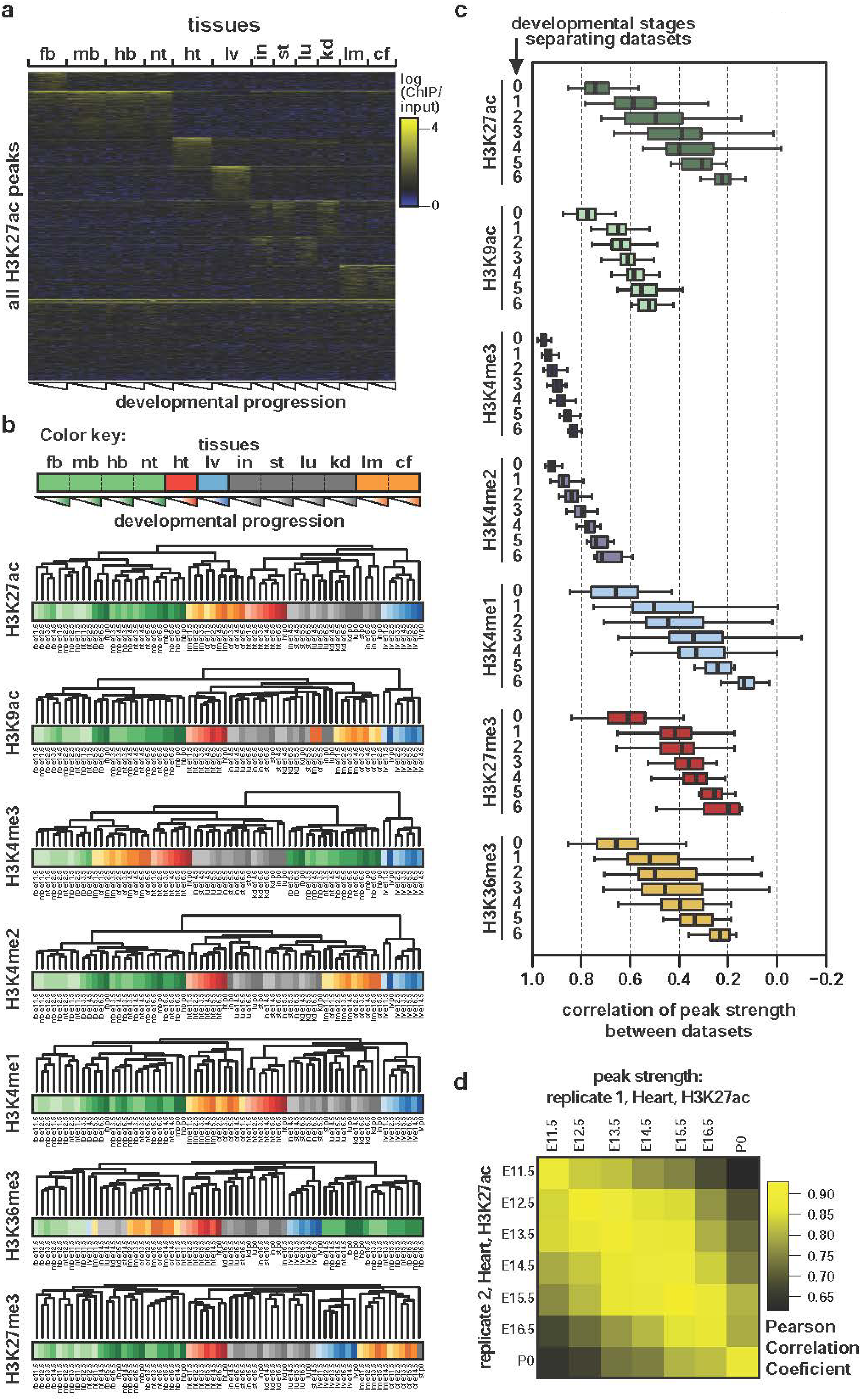
The histone modification landscape changes progressively during development and reflects developmental relationships between tissues. (a) k-means clustering of H3K27ac peaks based on normalized H3K27ac enrichment (see methods for details of normalization; k=8). Total number of H3K27ac peaks = 333,097. Numbers of elements in each cluster from top to bottom: 20497, 50790, 31043, 36849, 38670, 31168, 36822, 87258. Tissue abbreviations same as figure 1. (b) Dendrograms from hierarchical clustering based on signal for each histone modification, as indicated. Note the consistent relationships between tissues of similar developmental origin. (c) Progressive changes in histone modifications shown by pearson correlations of peak strength between replicates that are either from the same stage (i.e. developmental stages separating datasets = 0), or from different stages (separated by 1 - 6 intervening stages, as indicated). (d) Pearson correlation coefficients between all Heart H3K27ac datasets from E11.5 to P0.

### An integrated 15-state model characterizes the developmental chromatin landscape

To leverage the information captured by combinatorial patterns of histone modifications (i.e. chromatin states) we applied ChromHMM^13^ to the data series, excluding E10.5 because this stage did not have the full complement of histone modification profiles. We first determined the optimal number of chromatin states to describe the data (Figure 3a, Extended data figure 4a-c, and Methods). Multiple approaches converged on a 15-state model, which shows near-perfect consistency between biological replicates, and is in general agreement with previously published models developed from partially overlapping sets of histone modifications^13,16,24^ (Extended data figure 4d-g). Although our chromatin state designations were made without considering gene locations, the distribution of each state relative to genes matches expectations^25^ (Figure 3b). We grouped the resulting chromatin states into 4 broad functional classes: promoter states, enhancer states, transcriptional states, and heterochromatic states. It is important to note that these chromatin state annotations are specific to this study, and are distinct from the ENCODE Consortium’s candidate regulatory element registry and Encyclopedia (ENCODE Consortium, *in preparation*). In total, we find that ~33.4% of the genome shows a chromatin signature characteristic of one of these four functional classes (i.e. excluding state 15 “no signal”, and state 11 “permissive”) in at least one tissue or developmental stage. This does not necessarily mean that 33.4% of the genome sequence is functional, but rather that 33.4% of the genome sequence is packaged in chromatin that shows a reproducible signature in at least one of the tissues or stages profiled here. These chromatin signatures reflect transcriptional and/or regulatory activity, but the sequences they mark may not be functional by strict definition. For example, the transcriptional elongation state (state #10) covers the introns of many highly transcribed genes, but much of the sequence in these introns may not contribute to the fitness of the organism.

**Figure 3:**
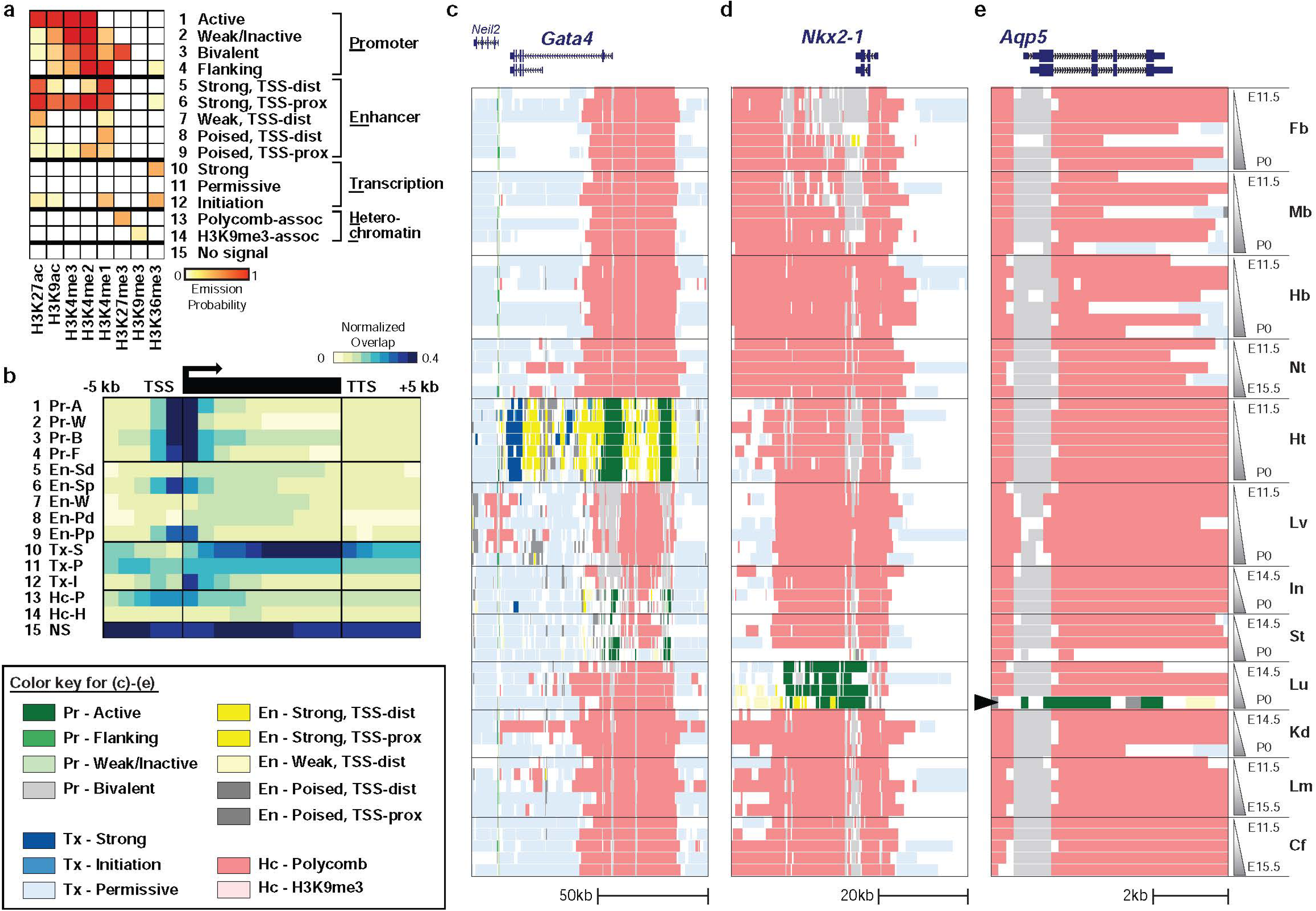
A 15 state model characterizes the mouse developmental chromatin landscape. (a) Overview of 15-state chromHMM model. A description is provided for each one of the 15 states. The states are grouped into four main categories: promoter, enhancer, transcription, heterochromatin. (b) Enrichment of chromatin states relative to annotated genes. The locations of these features were not considered when building the chromatin state model. (c) Chromatin state landscape at *Gata4*, a TF involved in heart development (chr14:63,181,234-63,288,624; mm10). (d) Chromatin state landscape at *Nkx2-1*, a TF involved in lung development (chr12:56,507,647-56,560,509; mm10). (e) Chromatin state landscape at *Aqp5*, a marker of mature Alveolar Type 1 cells in the lung, which is expressed just prior to birth (chr15:99,589,794-99,596,121; mm10). P0 lung stage indicated by black arrowhead.

The chromatin state maps provide powerful tools for genome annotation, as a broad range of inferred functional information can be visualized across multiple tissues and stages in a single genome browser view (Figure 3c-e; Extended data figure 5a-b). For example, we see that active chromatin states robustly and specifically mark master regulators of tissue development including *Gata4* and *Nkx2-5* in heart^26^, *Nkx2-1* in lung^27^, *Cdx2* in intestine^28^, *Barx1* in stomach^29^, and *Wt1* in kidney^30^. While examining chromatin state maps at these genes that code for tissue-restricted transcription factors (TFs), we noted that they are frequently marked by a heterochromatic state characteristic of Polycomb-mediated repression (state 13, Hc - Polycomb associated) outside of their tissues of activity. Polycomb-mediated repression is known to be essential for embryogenesis^31-33^, but the genomic distribution of repressive chromatin during mammalian development has not been thoroughly examined. To further investigate the role of Polycomb-mediated repression in development, we used GREAT^34^ to examine Gene Ontology (GO) terms associated with regions of Polycomb repressive chromatin. We found that Polycomb repressive chromatin is highly enriched near TF genes in every tissue and stage surveyed here (Figure 4a). Moreover, TF genes that are underlie human Mendelian diseases^35^ are particularly likely to be marked by repressive chromatin (Figure 4b), suggesting that these genes are tightly regulated by a combination of repression and activation. We also examined a small but well-characterized set of genes (*Sox9, Pax3, Shh, Ihh*, and *Wnt6*) that are known to cause human congenital phenotypes when ectopically expressed during development^36-38^. Notably, every one of these genes is widely marked by repressive chromatin in tissues where they are not active (Figure 4b; Extended data figure 5c). These data support an essential and pervasive role for Polycomb-mediated repression in silencing key developmental regulators outside of their normal expression domains, which may function in turn to control broader programs of gene expression^39^. We note that the heterochromatic state characterized by H3K9me3 (state #14, Hc - H3K9me3) and likely representative of facultative heterochromatin shows a very different distribution, as previously described^40-44^ (Extended data figure 6).

**Figure 4:**
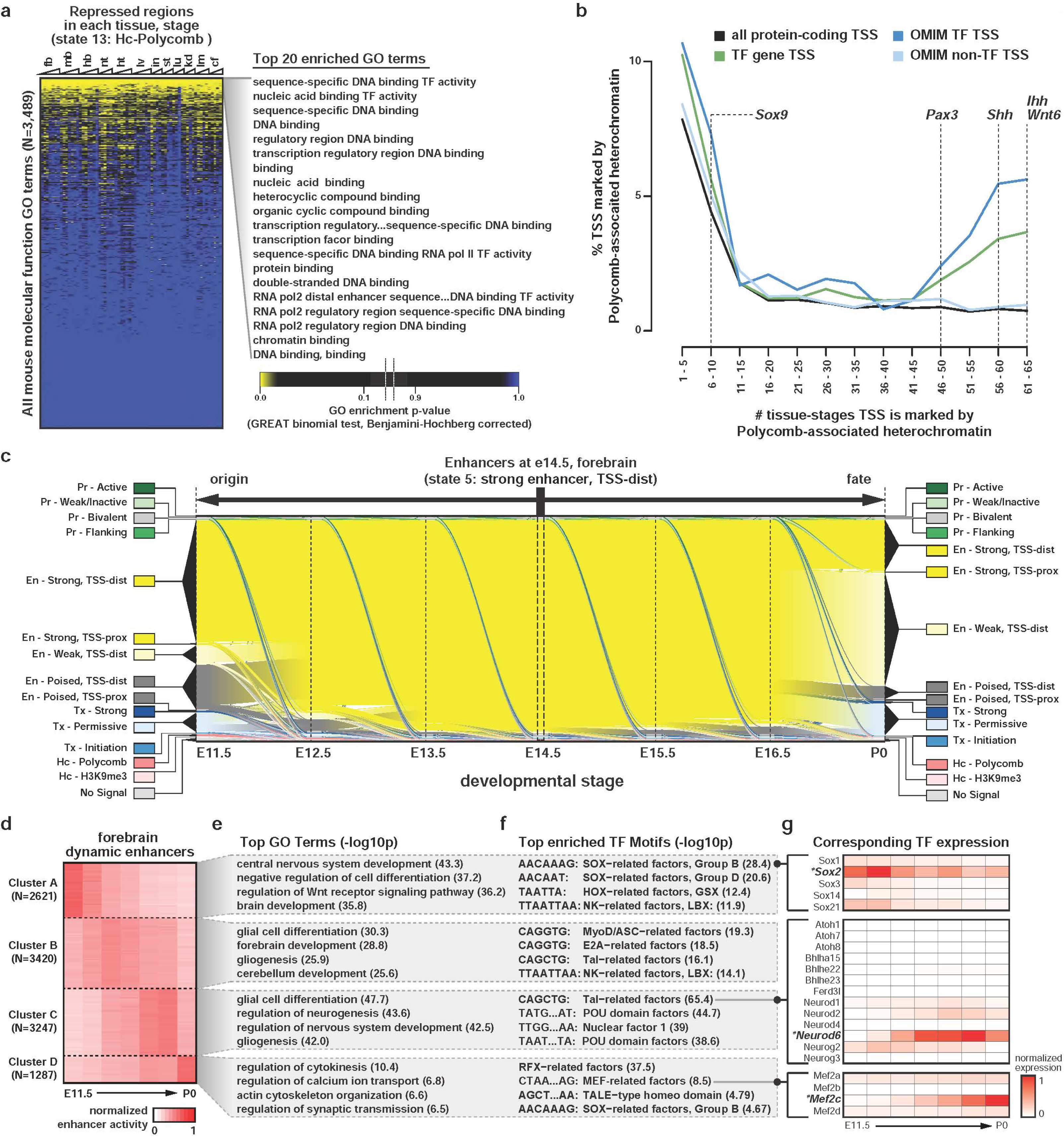
Repression and activation of cis-regulatory elements during development. (a) Enrichment of “molecular function” GO terms in genes near repressed regions (state 13). GO terms on the y-axis are ordered by average enrichment p-value across all tissues and stages (most significant on top to least significant on the bottom). The top 20 GO terms are listed on the right, and are all related to TF function. (b) The relative frequency of repressive chromatin (state 13) at four different sets of TSS: 1) all GENCODE protein coding TSS (N=90,384), 2) the subset of (1) that belong to transcripts of TF genes (N=1,019), 3) the subset of (2) that are also known to cause to Mendelian phenotypes when mutated (i.e. “OMIM” genes; N=227), 4) the remaining OMIM gene TSS that do not encode TFs (N=3,497). The individual gene symbols indicate genes that are known to cause congenital defects when abnormally expressed. Not shown: TSS repressed in none of the 66 tissue-stages (77%, 62%, 51%, and 74%, respectively), and TSS repressed in all 66 tissue stages (<1%, 2%, 3%, and <1%, respectively). (c) Sankey diagram showing the origin and fate of all genomic intervals classified as TSS-distal strong enhancers (state #5) in E14.5 forebrain. The chromatin state classification of these regions was tracked across the available developmental stages, and the relative genomic coverage of each chromatin state at each transition is plotted. The thickness of each color (i.e. y axis) indicates the coverage of each state. (d) K-means clustering of dynamic enhancers based on H3K27ac signal at stages: E11.5, E12.5, e13.4, E14.5, E15.5, E16.5, P0. (e) Top 4 Biological Processes GO terms enriched in each cluster of dynamic enhancers in (d) as determined by GREAT. (f) Top 4 sequence motifs enriched in each cluster of dynamic enhancers in (d). Some motifs are abbreviated with “…” to fit in the panel. The full motifs are (top to bottom): POU domain factors TATGCAAAT; Nuclear Factor 1 TTGGCTATATGCCAA; POU domain factors TAATTAATTA; MEF-related factors CTAAAAATAG; TALE-type home domain AGCTGTCAA. (g) Gene expression shown for TFs in the structural subfamilies that match motifs in (f) indicated by closed black circles.

The breadth of data collected here allowed us to globally characterize both the spatial and temporal dynamics of chromatin states. On average, ~1.2% of the genome differs in chromatin state between tissues at the same stage (mean 1.2%, 31.3Mb; range 1.0%-4.0%, 26.8Mb-109.1Mb), but that percentage jumps to 17.2% if we discard the portion of the genome that shows no chromatin signal across all tissue-stages (i.e. constitutive state 15). Enhancer states show the highest degree of variability between tissues, reinforcing the role of enhancers in defining tissue and cell identity (Extended data figure 7a). Indeed, hierarchical clustering of our samples based on strong enhancer predictions alone (i.e. state 5) leads to clear separation by tissue, and captures relationships between tissues with similar developmental origin (Extended data figure 7b-d). Interestingly, this analysis revealed a strong relationship between limb and facial tissue, which was also observed in clustering results based on single histone modifications (Figure 2a-b). These findings are suggestive of a deep similarity in gene regulatory programs between these two tissues, and provide further support for the hypothesis of a common developmental origin for facial structures and limbs^45,46^. Within a given tissue, we find that on average ~1.3% of the genome differs in chromatin state between adjacent developmental stages (mean 1.3%, 36.6Mb; range 0.03%-3.01%, 9.4MB-82.1Mb). While the fraction of the genome that differs in state between adjacent stages is roughly the same as the fraction that differs between tissues at the same stage, differences between stages are more subtle, as they are less likely to affect states characteristic of strong enhancers, promoters, or heterochromatin (States 5, 1, and 13, respectively; Extended data figure 7e). Nonetheless, these temporal changes capture important developmental processes like the maturation of Alveolar Type I cells (marked by *Aqp5^27^*) just prior to the onset of pulmonary gas exchange at birth (Figure 3e), and the transition of predominant liver function from hematopoiesis in the early embryo to metabolism in the later stages of development^47^ (Extended data figure 7f). A global summary of temporal changes in chromatin state reveals the tendencies of certain states to transition to others, including shifts from poised to active enhancers, bivalent to active promoters, and permissive to repressed chromatin (Figure 4c, Extended data figure 7g).

To gain further insight into the dynamic processes occurring in each tissue, we focused specifically on the subset of enhancers that show dynamic activity across stages (referred to as “dynamic enhancers” below). To identify dynamic enhancers, we began by compiling lists of putative enhancers in each tissue (i.e. chromatin state 5). Next, we calculated a stage-specific activity score for each enhancer based on its level of H3K27ac enrichment, and used these activity scores to identify enhancers that change activity from stage-to-stage^48^ (see **methods**). Within each tissue, clustering of these dynamic enhancers revealed several predominant temporal patterns of activity (Extended Data figure 8). Considering forebrain as an example, we found four predominant patterns of temporal activity, which reflect underlying developmental processes (Figure 4d-g). Enhancers that peak in activity early in development are associated with GO terms related to general CNS and brain development (Figure 4d, cluster A), while enhancers that peak in middle stages are associated more specifically with neurogenesis and gliogenesis (Figure 4d, clusters B-C), and enhancers peaking latest in the developmental series are associated with synaptic function (Figure 4d, cluster D). By comparing the TF binding motifs enriched in each cluster to the expression levels of TF genes that target these motifs, we identified specific TFs that likely regulate the biological processes revealed by GO analysis. Consistent with published reports, these data suggest key roles for Sox2 in early brain development^49,50^, NeuroD6 in neurogenesis during mid-late gestration^51^, and Mef2c in synapse formation^52^. Notably, the human orthologs of *Mef2c* and *Sox2* are Mendelian disease genes linked to congenital defects of the CNS (MIM # 613443 and #206900, respectively). Similar catalogs for each tissue are available in Extended data figure 8 and Table S1, including dynamic enhancer clusters, corresponding GO terms, and enriched TFs.

### Linking Enhancers to Target Genes and Phenotypes

Predicting the target genes of enhancers remains a major challenge in the field, in part because enhancers can be located hundreds of kilobases away from the genes they regulate^22,23^. To address this challenge, we developed a map of predicted enhancer target genes based on the correlation between gene expression (as measured by RNA-seq) and H3K27ac enrichment at TSS-distal enhancers within the same Topologically-Associating Domain (TAD) (Figure 5a; Extended data figure 9a-b; Table S2). TADs have been shown to constrain the activity of enhancers to the genes in the same domain^36,37,53^. We assigned enhancers to target genes within the same TAD based on <u>S</u>pearman’s rank Correlation Coefficient (*SCC*) between H3K27ac enrichment and mRNA expression across all 66 tissue-stages (Extended data figure 9a) (*p*-value <= 0.05 and *SCC* >= 0.25; see methods). We took advantage of biological replicates for both the chromatin and RNA-seq data to derive independent target gene maps for each replicate. These maps include 31,965 and 32,735 enhancer-gene assignments, respectively, with an overlap 21,142 (Extended data figure 9c-e). Overall, these target gene maps are highly reproducible (SCC = 0.71 between replicates, evaluated as the number of enhancers per gene). Interestingly, we found that genes with 5 or more assigned enhancers are associated with biological processes related to development and differentiation, whereas genes with only a single assigned enhancer tend to be involved in cellular and metabolic processes more consistent with “housekeeping” functions (Extended data figure 9f). This result further highlights the importance of enhancers in directing the expression of developmental genes.

**Figure 5:**
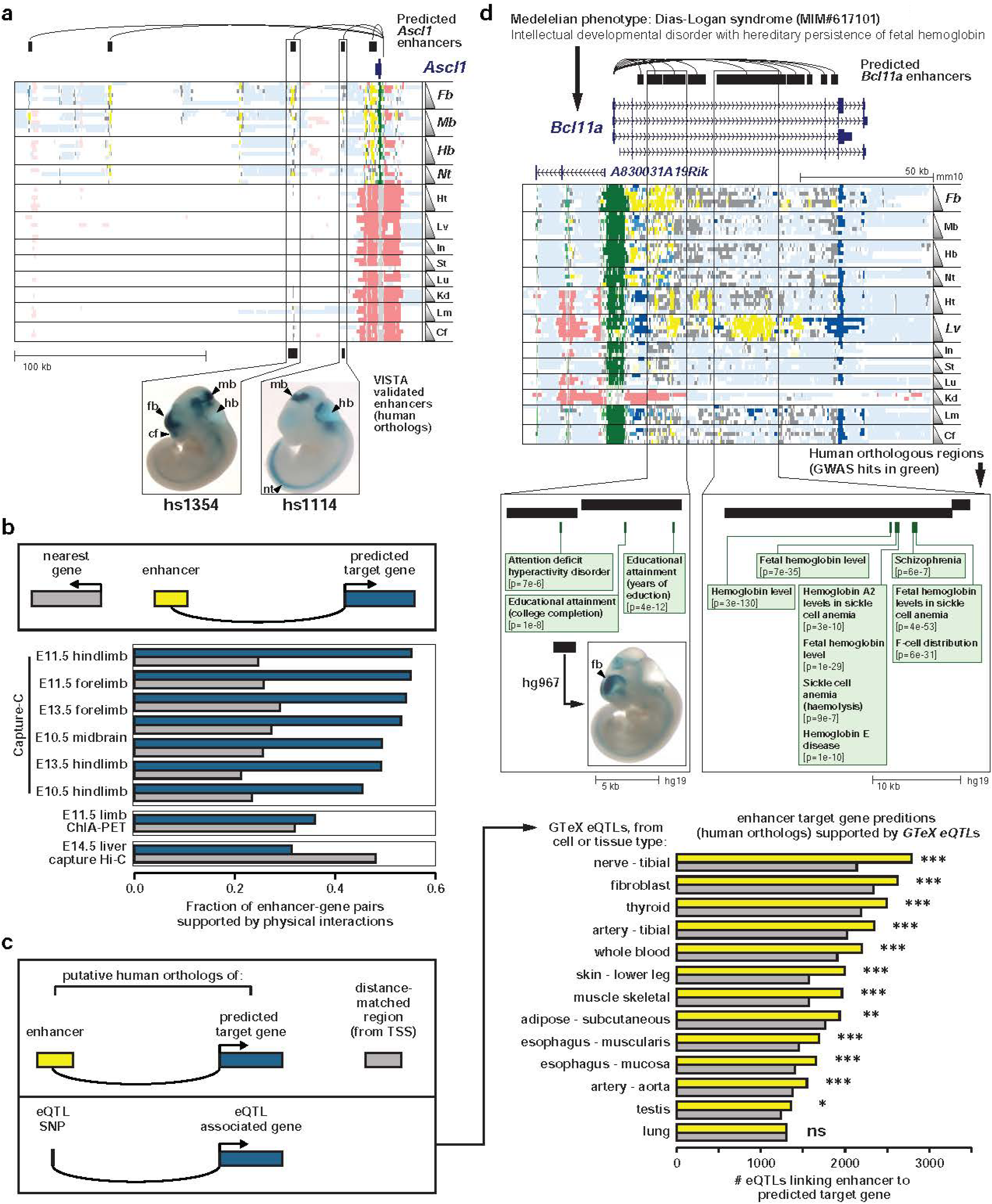
Maps of enhancer target gene prediction in mouse and human. (a) Genome browser view showing chromatin states at the *Ascl1* locus (chr10:87,301,848-87,515,210; mm10). Color scheme is the same is in Figure 4. Black bars on top represent enhancers predicted to target *Ascl1*. Two of these enhancers have human orthologs (below main panel) that have been previously validates using *in vivo* transgenic reporter assays in mouse. Tissues with reproducible enhancer activity are indicated with arrow heads. (b) Enhancer-gene interactions identified by this correlative approach are generally more likely to be supported by chromatin interaction data than associations derived by a nearest gene approach. Chromatin interaction data from (Andrey et al., 2017) (upper box), (DeMare et al., 2013) (middle box) and (Schoenfelder et al., 2015) (lower box). Note that the Liver capture Hi-C dataset contains ~600,000 interactions (or roughly 30 per TSS considering ~22K TSS in the genome). This is far more interactions than that Capture C or ChIA-PET datasets, which we expect may explain why the nearest gene assumption outperforms our predictions in this dataset alone. (c) The correlative enhancer-gene interactions are significantly enriched for human eQTL-gene associations (see Methods). *** p <= 0.001; ** 0.001 < p <= 0.01;* 0.01 < p <= 0.05; (Fisher’s Exact Test). (d) Genome browser view showing chromatin states at the *Bcl11a* locus (chr11:24,044,043-24,197,927; mm10). Boxes outline enhancer clusters active in the CNS (left) and liver (right). The subpanels below show regions of the human genome corresponding to the CNS enhancer cluster (chr2:60,752,530-60,767,198; hg19) and liver enhancer cluster (chr2:60,711,940-60,741,118; hg19), respectively. The black bars on top represent the predicted Bcl11a enhancers from mouse, and green bars below represent GWAS SNPs for the EMBL-EBI GWAS catalog. SNP IDs lower left subpanel, from left to right: (rs2556378, rs2457660, rs356992). SNP IDs lower right panel, left to right: (rs4671393, rs11886868, rs766432, rs7599488, rs1427407). The black bar labeled hs967 shows a sequence previously tested *in vivo*. Reporter activity in E11.5 mouse embryo shown to the right.

To further evaluate the accuracy of our enhancer target gene maps, we made use of published data from chromatin conformation capture technologies^54-56^, which assay physical interactions between promoters and enhancers. We found that our correlation-based map predicts experimentally determined enhancer-gene interactions with higher accuracy than assigning an enhancer to the nearest adjacent gene (Figure 5b; Extended data figure 9g). We also sought to determine whether our map of enhancer target genes in mouse could be useful for predicting human enhancer-gene interactions. To this end, we identified human orthologs of the enhancers and genes in our mouse map, and used data from the GTEx project to assess whether the enhancer-gene relationships inferred in mouse are likely to be conserved in human. The GTEx project has identified a large collection of human expression quantitative trait loci (eQTLs)^57^, which are known to be enriched in enhancers and other regulatory elements^58^. Importantly, each eQTL is connected to a target gene by genetic association. Thus, we hypothesized that if the enhancer target genes we predict in mouse are applicable to human, we should see an enrichment for eQTLs that link the human orthologs of these enhancers to the same target genes by genetic association. Indeed, across a variety of human tissues we see a significant enrichment of eQTLs that support our enhancer target gene predictions (Figure 5c).

With this set of enhancer target gene predictions in hand, we again focused on the subset of genes known to cause Mendelian disease when mutated (“OMIM genes”)^35^. Mendelian diseases are predominantly caused by highly penetrant mutations that often effect coding sequence^59^. In contrast, SNPs associated with lower-penetrance non-Mendelian phenotypes by Genome-Wide Association Studies (GWAS) are highly enriched in putative regulatory sequences^60^. This suggests that dysregulation of gene expression is a common mechanism contributing to non-Mendelian phenotypes, which include diseases with enormous human health burden like coronary heart disease^61^ and Type 2 diabetes^62^. We hypothesized that if enhancers which regulate Mendelian disease genes are disrupted by common sequence variants (i.e. SNPs), it might lead to less penetrant non-Mendelian phenotypes. To explore this hypothesis, we examined the frequency of GWAS SNPs (i.e. SNPs associated with a human phenotype by GWAS) in enhancers linked to OMIM genes by our correlative map, and compared this with the frequency of GWAS SNPs in enhancers linked to non-OMIM genes. Indeed, we find a highly significant enrichment for GWAS SNPs in enhancers linked to OMIM genes (*p*=4.3e-8, chi-squared test). We also compared enhancers linked to OMIM genes with enhancers in the same TADs that are not linked to OMIM genes. Again, we see a significant enrichment for GWAS SNPs in the enhancers linked to OMIM genes (p=7.29e-4, chi-squared test; Extended data figure 9h; Supplementary Table S4). These results points to a systematic relationship between highly penetrant mutations that lead to Mendelian disease, and lower penetrance enhancer variants that may impact the regulation of those same genes and lead to more common phenotypes. For example, in Figure 5d we highlight *Bcl11a*. Coding mutations in the human ortholog (*BCL11A*) cause Dias-Logan syndrome (MIM# 617101) – also known as “Intellectual developmental disorder with hereditary persistence of fetal hemoglobin.” The *Bcl11a* locus contains two large clusters of putative enhancers that are active in the CNS and embryonic liver (a site of erythropoiesis), respectively. Indeed, the human ortholog of the liver enhancer cluster has been extensively characterized in human blood lineages^63,64^, and a human sequence orthologous to the CNS enhancer cluster has been validated *in vivo*65. Interestingly, the human liver enhancer ortholog contains nine total GWAS associations, eight of which are related to hemoglobin levels (the 9^th^ association is to schizophrenia). In contrast, the CNS enhancer orthologs contains three GWAS associations, all with phenotypes related to neurodevelopment or cognitive function (Attention deficit hyperactivity disorder, Educational attainment). These observations suggest that while *Bcl11a* coding mutations lead to a pleiotropic syndrome with manifestations in both the CNS and hematopoietic systems, non-coding regulatory variants at the same locus may lead to nonsyndromic phenotypes that predominantly effect one system or the other, depending on which enhancer is affected.

### Relationship between H3K27ac enrichment and enhancer validation rate

In several of the analyses described above, we have used chromatin state to study putative enhancers. Chromatin states (and even H3K27ac alone) have proven to be effective tools for identifying enhancers^14,48,66,67^, but the accuracy of these methods remains under investigation. The putative enhancers identified in this study are supported by multiple lines of evidence including the GO and motif enrichments discussed above (Figure 4f) as well as tissue-specific enrichments for *in vivo* validated enhancers from the VISTA database^65^ (Extended data figure 10a). Anecdotally, we have noted that putative enhancers with stronger H3K27ac enrichment are more likely to validate in transgenic mouse reporter assays, which is an important consideration because the level of H3K27ac enrichment at putative enhancers can vary by orders of magnitude within a single tissue or cell type. To more systematically examine this relationship between H3K27ac enrichment level and validation rate, we used transgenic mouse assays^68^ in E12.5 embryos to test 150 enhancers predicted from E12.5 forebrain, heart, or limb tissue that fell into one of three H3K27ac enrichment rank categories: high (selected from ranks 1-85), medium (ranks 1500-1550), or low (ranks 3000-3050) (Figure 6, Extended data figure 10b, Supplementary Table S5). We found that ~60% of the high-ranking enhancers displayed reproducible reporter gene expression in the expected tissue. However, <30% of predicted enhancers from the two lower-rank groups validated (Figure 5a, p-values < 0.01, Fisher’s exact test). These results are consistent with a similar validation of enhancers predicted from e11.5 tissues (ENCODE Consortium, *in preparation*). In addition, high-ranking enhancers that validated in the expected tissue were also more likely to have enhancer activity in additional tissues (Figure 6b, p-values < 0.05, Mann-Whitney U Test). Importantly, no significant differences were observed in reproducibility rates (i.e. the percentage of embryos with reproducible activity) between rank classes (Extended data figure 10c). Overall, these results demonstrate that loci with stronger H3K27ac enrichment are more likely to direct reporter expression in the expected tissue, and tend to direct reporter expression in a wider variety of tissues.

**Figure 6:**
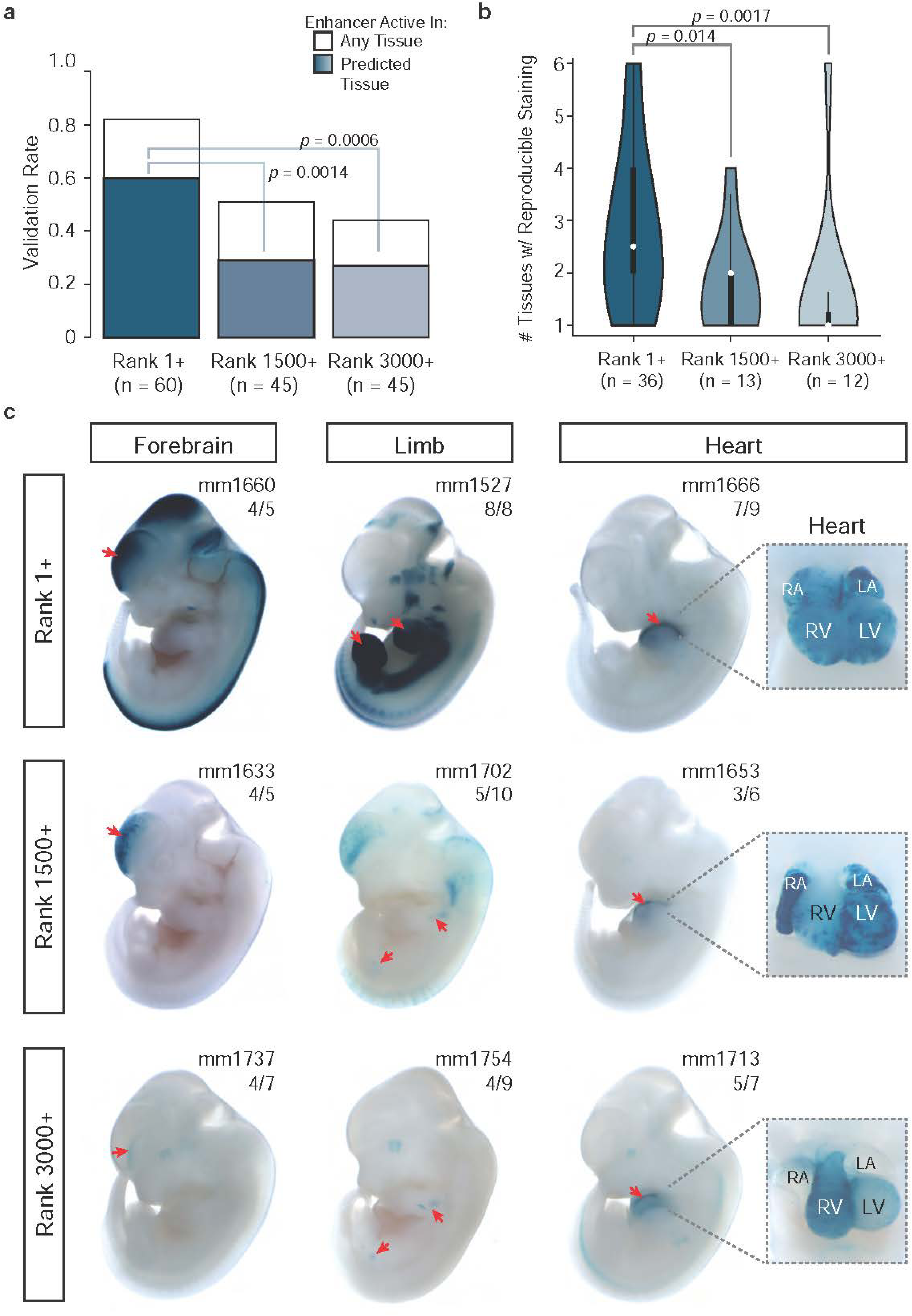
*In vivo* validation of predicted E12.5 enhancers. We tested 150 total enhancers predicted from E12.5 forebrain, limb, or heart tissue using in vivo mouse transgenic assays. Predicted enhancers from each tissue were ranked, from strongest to weakest H3K27ac signal, and we selected elements for validation from three different rank categories (roughly ranks 1-85, 1500-1550, and 3000-3050). (a) Bars indicate the proportion of enhancers in each category that showed reproducible reporter staining in the expected tissue (blue) or any tissue (white). High ranking predicted enhancers had a higher validation rate in the correct tissue than lower ranking enhancers (p-values by 1-tailed Fisher’s exact test). (b) For those enhancers that validated in the expected tissue, the distribution of tissues with reproducible reporter gene expression for each enhancer by rank class is shown with violin plots. High ranking enhancers showed reproducible activity in more tissues than those with lower ranks (p-values by one-tailed Mann-Whitney U Test). (c) Examples of enhancers from each tissue type and rank class that validated in the expected tissue. Representative transgenic E12.5 embryos showing reporter expression (blue staining) are shown, along with the unique VISTA Enhancer Browser (PMID: 17130149) identifier (mm number) and the enhancer’s reproducibility. Images on the far right show a magnified image of the heart (RA, right atrium; LA, left atrium; RV, right ventricle; LV, left ventricle). Red arrowheads indicate the enhancer activity pattern in the forebrain, limb, or heart.

The results of these transgenic reporter assays provide an intriguing perspective on one of the most important questions in the field: what portion of sequences marked by an enhancer chromatin signature truly function as enhancers *in vivo*? Surveys of chromatin state and chromatin accessibility in a single tissue or cell type often predict enhancers numbering in the tens or even hundreds of thousands. In the past this has led us and others to estimate the total number of enhancers in the genome at several hundred thousand or more^66,69^. In contrast, the results of our transgenic assays suggest that outside of the top enhancers in a given tissue (as ranked by H3K27ac enrichment), the validation rate in this assay drops to roughly one-third. In light of these results, we think it is important to approach estimates of enhancer abundance with a healthy degree of caution, but we do not think these estimates should be abandoned entirely as they likely provide a reasonable upper bound estimate for the number enhancers in the genome. Although our data clearly suggests that the true positive rate of chromatin-based enhancer predictions decreases as function of H3K27ac level, it is difficult to determine exactly what this true positive rate is. Definitive proof of an enhancer’s function (or lack thereof) requires genetic manipulation of that enhancer in its native context, followed by extensive measurements of the resulting impact on gene expression^70-72^. Even such rigorous genetic experiments can be difficult to interpret, because an enhancer’s function might be limited to a very specific set of conditions that are difficult to identify and/or reproduce in an experimental setting. In addition, one enhancer may compensate for the loss of another, achieving a redundancy that could serve to make regulatory programs more robust to perturbations. Ultimately, we think our results highlight the importance of continued investigation into the molecular basis of enhancer function, and encourage the continual assessment of global enhancer estimates as new data points like those reported here become available.

In summary, we present a survey of the chromatin landscape in the developing mouse embryo that is unprecedented in is breadth. Our results describe a multi-tiered compendium of functional annotations for the developmental mouse genome, yielding valuable insight into key developmental processes and regulatory factors. The resources generated here include chromatin state maps for each tissue and stage, catalogs of enhancers that show dynamic activity during development, a genome-wide map of predicted enhancer target genes, and a collection of transgenic reporter assays that demonstrates a strong relationship between H3K27ac signal and likelihood of validation. We further reveal an important role for Polycomb-mediated repression in regulating TF genes during development, and a systematic connection between Mendelian diseases and complex common diseases likely caused by perturbation of the same genes. Due to the uniquely critical role of the mouse in biomedical research, we believe that the tools and insights developed here will prove to be a valuable resource to the biomedical research community.

## AUTHOR CONTRIBUTIONS

Study was conceived and overseen by B.R., D.U.G., L.A.P., A.V., D.E.D., Y.S., F.Y., J.E., W.W, and J.M.C. Tissue collections performed by V.A., J.A.A., I.P-F., C.S.P., and M.K. ChIP-seq experiments performed by A.Y.L. and S.C. Data curation and processing by J.S.S., J.M.D., and D.U.G. Data analysis performed by D.U.G., I.B., Y.F-Y., Y.Zhang, A.W., B.D., B.Z., Y. Zhao, M.W., Y.Q., Y.H., and S.P. Transgenic assays performed by V.A., J.A.A., I.P-F., C.S.P., M.K., T.H.G., Q.T.P., A.N.H., B.J.M., and E.A.L. Manuscript written by D.U.G., I.B., D.E.D., and Y.X. with input from all authors.

## ACKNOWLEDGMENTS

This study was funded by the National Human Genome Research Institute as part of the Encyclopedia of DNA Elements (ENCODE) project (U54HG006997). D.U.G. was supported by the NIH Institutional Research and Academic Career Development Awards (IRACDA) program, and a postdoctoral fellowship from the A.P. Giannini Foundation. D.E.D, A.V., and L.A.P. were also supported by NIDCR FaceBase grant U01DE024427, NHLBI grant R24HL123879, and NHGRI grants R01HG003988, and UM1HG009421, and research conducted at the E.O. Lawrence Berkeley National Laboratory was performed under Department of Energy Contract DE-AC02-05CH11231, University of California. Y.H. is supported by the H.A. and Mary K. Chapman Charitable Trust. J.R.E. is an Investigator of the Howard Hughes Medical Institute.

## METHODS

### Tissue Collection

All animal work was reviewed and approved by the Lawrence Berkeley National Laboratory Animal Welfare and Research Committee. Tissue collection for all developmental stages was performed using C57BL/6N strain *Mus musculus* animals. For E14.5 and P0, breeding animals were purchased from both Charles River Laboratories (C57BL/6NCrl strain) and Taconic Biosciences (C57BL/6NTac strain). For all remaining developmental stages, breeding animals were purchased exclusively from Charles River Laboratories (C57BL/6NCrl strain). Wild-type male and female mice were mated using a standard timed breeding strategy. Embryos and P0 pups were collected for dissection using approved institutional protocols. Embryos were excluded if they were not the expected developmental stage. To avoid sample degradation, only one embryonic litter or P0 pup was processed at a time, and tissue was kept ice cold during dissection. Collection tubes for each tissue type were placed in a dry ice ethanol bath so that tissue samples could be flash frozen immediately upon dissection. Tissue from multiple embryos was pooled together in the same collection tube, and at least two separate collection tubes were collected for each tissue-stage for biological replication. Tissue was stored in a −80°C freezer or on dry ice until further processing. A step-by-step protocol for tissue collection, including detailed information about how embryonic stage was determined, can be found on the ENCODE Project website at https://www.encodeproject.org/documents/631aa21c-8e48-467e-8cacd40c875b3913/@@download/attachment/StandardTissueExcisionProtocol_02132017.pdf.

### ChIP-seq data generation

ChIP-seq experiments for all marks and tissues from E11.5 – P0 were performed as previously described^66^. The ChIP-seq protocol was modified slightly for all E10.5 experiments due to the low amount of input (“micro” ChIP-seq). Detailed protocols for both standard and micro ChIP-seq including antibodies used and antibodies validations performed are available at www.encodedcc.org associated with each experiment described here. They can also be found at the links below. Standard ChIP-seq, Tissue fixation & sonication: https://www.encodeproject.org/documents/3125496b-c833-4414-bf5f-84dd633eb30d/@@download/attachment/Ren_Tissue_Fixation_and_Sonication_v060614.pdf, Immunoprecipitation: https://www.encodeproject.org/documents/89795b31-e65a-42ca-9d7bd75196f6f4b3/@@download/attachment/Ren%20Lab%20ENCODE%20Chromatin%20Immunoprecipitation%20Protocol_V2.pdf, Library preparation: https://www.encodeproject.org/documents/4f73fbc3-956e-47ae-aa2d-41a7df552c81/@@download/attachment/Ren_ChIP_Library_Preparation_v060614.pdf. Micro-ChIP-seq (E10.5), Tissue fixation & sonication: https://www.encodeproject.org/documents/1fcaab50-6ca0-4778-88cb-5f6b85170d21/@@download/attachment/Ren%20Lab%20ENCODE%20Tissue%20Fixation%20and%20Sonication%20Protocol%20MicroChIP.pdf, Immunoprecipitation & Library preparation: https://www.encodeproject.org/documents/18580e80-0907-4258-a412-46bcc37bd040/@@download/attachment/Ren%20Lab%20ENCODE%20Chromatin%20Immunoprecipitation%20Protocol%20MicroChIP.pdf.

### ChIP-seq data processing

Histone ChIP-seq data were analyzed using a software pipeline implemented by the ENCODE Data Coordinating Center (DCC) for the ENCODE Consortium. A schematic representation of the pipeline is shown in Extended figure 10. Each step of the pipeline corresponds to a script written in the Python programming language that assembles the input files, runs external programs (such as the MACS2 peak caller), and calculates quality-control metrics. The methodology is similar to that previously described for ENCODE^73^ with the following modifications: The mapping step used bwa version 0.7.10 and samtools version 1.0, and MACS2 version 2.1.0 was used for signal track generation and peak calling. In order to ensure an adequate sampling of noise for subsequent replicate comparisons, peaks were initially called at a relaxed p-value threshold of 1 x 10-2. Such relaxed peak sets were generated for each biological replicate, for the replicates pooled, and for pooled pseudoreplicates of each true replicate (each pseudoreplicate consists of half the reads sampled without replacement). Peaks from the pooled replicate set were retained in the replicated peak set if they overlapped by at least half their length (in bases) peaks from both biological replicates. Additionally, peaks that overlapped both pooled pseudoreplicates were added to the replicated peak set. In this way very strong biological replicates could “rescue” peaks that were only marginal in a second replicate. The pipeline is available to be run on the DNAnexus (https://www.dnanexus.com/) web platform, backed by cloud computing from Amazon Web Services (AWS), and is the same pipeline used for the analysis of all ENCODE histone ChIP-seq experiments. The platform provides both an API for programmatic execution of the pipeline and a web-based interface for interactive execution of the same workflows. ENCODE DCC uses this approach to ensure that primary data from different labs within the Consortium are processed uniformly, and thus minimizing factors that could confound subsequent comparisons^74^. The ENCODE DCC analyzed the experiments in parallel and accessioned the results to the ENCODE Portal^73^ (https://www.encodeproject.org/).

### Code availability

The ENCODE histone ChIP-seq pipeline is among the collection of ENCODE Uniform Processing Pipelines that can be found here: https://platform.dnanexus.com/projects/featured. The code is open-source, and available here: https://github.com/ENCODE-DCC/chip-seq-pipeline.

### Analysis of individual histone ChIP-seq data. K means clustering

To facilitate clustering across all tissues and stages, one master H3K27ac peak list was generated by merging peaks across tissue-stages. Each peak was then scored for H3K27ac ChIP-seq fold enrichment over input in each tissue-stage using *bigWigAverageOverBed*, and these values were quantile normalized across tissue-stages to eliminate potential confounding effects of systematic biases in the distribution of signal between tissues and/or stages. These normalized values are represented by the blue-black-yellow color scale in Figure 2a. One additional data processing step was performed prior to clustering: across each row, the values were converted to a unit vector in R (x / sqrt(sum(x^2)), to prevent enrichment level within tissue-stages from dominating the clusters, rather than enrichment level between tissue-stages. Note that these unit vector values were only used for clustering; the values plotted are the normalized H3K27ac enrichment values. K means clustering was performed in R with k = 8 and default parameters. Rows were ordered within each cluster based on mean normalized enrichment. **Hierarchical clustering of tissue-stages**. Normalized ChIP-seq fold enrichment values were calculated for each mark as described for H3K27ac in the preceding subsection (not including unit vector conversion). Hierarchical clustering was performed in R with default parameters. The resulting dendrograms are plotted in Figure 2b. **Calculation of distance between stages**. For each mark, we used Pearson correlation to compare the similarity between all datasets from the same tissue. We then categorized these correlations values based on how many stages separate the datasets being compared: 0 for true biological replicates from the same stage up to 7 for comparisons of E11.5 datasets to P0 datasets. These correlations are plotted in Figure 2c. **Principle Component Analysis (PCA)**. The whole genome was split into 1kb tiling bins. Average fold enrichment signals were calculated for each bin using the bigwigoverbed.sh script from the UCSC website. Bins that overlapped a merged peak by a minimal 20% (reciprocal) was denoted as peak-bin. The average fold enrichment signals from each peak-bin were quantile normalized within the tissue (not across the tissue). The signal strength for each peak was calculated as the sum of the signals of all bins overlapping that peak. Principal component analysis was performed on the peak signals for each histone mark with the R function “prcomp”. PC1, PC2 and PC3 values were plotted for each sample. **Metagene profiles**. To illustrate the characteristic enrichment patterns at active and silent genes, we used very conservative definitions to define these gene sets here. Active genes are defined as RPKM > 10 in every tissue-stage evaluated here (see Table S6 for list of public RNA-seq datasets). Silent genes were defined as RPKM < 2 in all tissue-stages. Metagene profiles were plotted with deeptools plotProfile^75^, using data from E15.5 Heart. **Correlation between replicates**. To facilitate comparisons of peak strength between replicates, we began with a single list of peaks called for each tissue-stage using data pooled from both replicates. Then, we scored these peaks for ChIP-seq fold enrichment over input for each replicate separately using *bigWigAvgOverBed*, and performed quantile normalization to eliminate confounding effect of systematic biases in the distribution of signal between replicates. We then used Pearson Correlation to compare the replicated of each experiment. “Narrow” marks (H3K4me3, H3K4me2, H3K27ac, H3K9ac) for tighter peaks of enrichment and tend to correlate more strongly than “broad” marks (H3K27me3, H3K4me1, H3K9me3, H3K36me3).

### ChromHMM. Generating the model

Chromatin datasets (bam files) were downloaded from the ENCODE DCC on October 15, 2016. De-duplicated bam files for each sample, along with their respective input controls, were binarized using the *binarizeBam* function of ChromHMM, with default parameters. Models considering 2 to 24 states were learned separately on the two replicates using the *LearnModel* function, with default parameters. For the rest of the analyses, we leveraged the availability of two, distinct time series; namely, we applied the same strategy separately and compared the results *a posteriori*. The conclusions obtained were invariably consistent, suggesting that the inferences on a single time series (at least in terms of global genomic patterns) are highly reproducible. **Identifying the optimal number of chromatin states**. We devised two different strategies to identify the minimal number of states that capture the combinations of histone modifications present in the data, both of which converged on a 15 state model. First, the ChromHMM *CompareModels* function was run separately on the two series. This function compares the emission parameters of a selected model to a set of models (in terms of Pearson’s correlation), and outputs the maximum correlation of each state in the selected model with its best matching state in each other model. We used this function to compare the “full” model (namely the one that consider 24 states) to the states in the simpler models. We then calculated the median correlation of all the 24 states against the simpler models, plotted these numbers against the number of states in the model and look at what number of states both series reached a plateau. As a complementary strategy, the emission probabilities from all the 23 models (considering 2 to 24 states) from both replicates were clustered together. The rationale behind this strategy is that very similar states across models will tend to cluster together, so there must be an optimal number of clusters corresponding to the optimal number of states in the model. To this end, we applied *k*-means clustering with *k* between 2 and 24, and evaluated the goodness of the separation for each *k* as the ratio between the “Between sum of squares” (referred to as Between SS) and the “Total sum of squares” (Total SS). Very cohesive, well-separated clusters tend to approach a ratio of 1. Given a value of *k*, the ratio was averaged over one hundred realizations of the clustering. The ratio observed for *k* = 24 was used as maximum, and the optimal number of states was then defined by the smallest value of *k* showing a ratio equal or higher than 95% of the maximum. **Genome segmentation and chromatin state tracking across genomic positions**. The segmentation was run separately for each sample, using the *MakeSegmentation* function of ChromHMM (default parameters) and the model derived from the first replicate. For the final set of replicated state calls we required that a region was assigned to the same state in both biological replicates (within a given tissue & stage). Regions that were not assigned to the same state in both replicates were re classified as “No Reproducible Signal” (distinct from state 15 – no signal in both replicates). The *unionbedg* functionality of BEDTools76 v2.17.0 was exploited to keep track of the chromatin state of genomic intervals across a defined set of samples. **Chromatin state trajectories along developmental time**. Given a replicate for a defined tissue, all those genomic intervals classified in a specified state (e.g. #5, strong enhancers) in one or more time points were tracked using the approach described above. Considering each pair of adjacent developmental time points, the genomic coverage of each transition between each pair of states was then calculated. The resulting numbers were then normalized on the coverage of the largest transition in the time series under investigation (e.g. liver, replicate #2) and shown as a directed graph. **Clustering on enhancer states**. After tracking the changes in chromatin state of each genomic base pair in the genome across multiple stages and tissues, the resulting matrix was binarized according to each segment being classified in a specified state (1) or any other state (0). The binary distances between all the pairs of samples considered in each specific analysis were then calculated. These were used either for comparisons or hierarchical clustering (Ward’s method). **GO analysis**. Functional enrichments through GREAT^34^ were obtained through the *greatBatchQuery.py* script. The resulting lists were first filtered for the relevant ontologies. After that, only the terms showing a binomial FDR <= 0.05 and a regional enrichment equal or higher than 2-fold were considered. WebGestalt^77^ was run with default parameters, specifically restricting the enrichment analysis to the GO Biological Process Ontology. VISTA validated elements were downloaded from https://enhancer.lbl.gov on June 17, 2016. Mm9 and hg19 coordinates were converted to mm10 using *liftOver* (setting *-minMatch* to 0.95 and 0.1, respectively). VISTA positive elements with any of the following annotations: "forebrain", "midbrain", "hindbrain", "neuraltube", "limb", "facialmesenchyme" or "heart" were considered for the following analysis. Liver was not considered in this enrichment analysis since there are currently <10 validated elements in VISTA showing reproducible staining in the liver. *coverageBed* from BEDTools v2.17.0 was used to calculate the coverage of the regions in each state in the E11.5 predictions with each tissue-specific group of VISTA elements. The fraction of bases covered was then normalized to the expected overlap, based on the overall genome-wide coverage of each state. The enrichment for repetitive elements was calculated using the *OverlapEnrichment* function of ChromHMM. **Classification of TSS as active, repressed, OMIM, and/or TF**. A 2kb window was defined around the TSS coordinates of all protein coding transcripts in GENCODE^78^ vM12. These 2kb windows (one per TSS) we overlapped with replicated chromHMM calls to determine their chromatin state in each tissue and stage. A TSS was classified as “active” in a given tissue-stage if this 2kb window overlapped the active promoter state (#1), and did not overlap any repressive states (#3, #13, #14). A TSS was classified as “repressed” in a given tissue-stage if this 2kb window overlapped a state characteristic of polycomb-mediated repression state (#3, #13), and did not overlap any active states (#1, #3, #4, #5, #6, #7, #10, #12). TSS that did not meet the criteria for either active or repressed in a given tissue-stage were left unclassified. Mouse orthologs of OMIM genes were taken directly from the “genemap.txt” available from www.omim.org, and filtered for genes with a “Disorder” type value of 3 – “the molecular basis of the disorder is known”. The full list of mouse TFs was downloaded from TFClass79. Ensembl Gene IDs were used to match GENCODE TSS to corresponding OMIM and TF genes. **Characterization of dynamic enhancer elements.** The temporal dynamic analysis was performed for each tissue separately. First, 1kb genomic bins that overlapped with regions defined as ChromHMM strong enhancer states in at least one stage were identified. Then we selected dynamic elements (bins) from these strong enhancer bins using the bioconductor LIMMA package^80^. LIMMA is a package developed for calling differentially expressed genes for microarray but was also adapted for sequencing data with the LOOM functionality. LIMMA package was used to call differential binding between each adjacent stage comparisons (e.g. E11.5 vs E12.5, E12.5 vs E13.5, etc). P-values were calculated with the eBayes function within LIMMA with trend parameter disabled, and were adjusted using Benjamini-Hochberg method. A bin was called overall dynamic if its adjusted p-value is less than 0.05 in any adjacent stage comparisons; otherwise it is called a non-dynamic bin. Non-dynamic bins were not included in the following analysis to reduce noise. We performed K-means clustering on dynamic bins across stages. The rows (bins) are normalized by dividing a common value so that the squares of the values sum up to 1. The optimal K was determined using the “elbow method” to cut off at the K value where percentage of withinness values transition from increasing quickly to increasing steadily with larger K. The resulting heatmap of the K-means clusters were shown in Figure 4d and Extended data figure 8. For each of the identified clusters, we performed enrichment testing of “GO Biological Processes” using GREAT. Over-represented motifs for each dynamic cluster was identified with the following method: First, all vertebrate Motif position weight matrices (PWMs) were downloaded from JASPAR TF database and used to scan the peak-bins for motif occurrences with FIMO, MEME suite^81^. For each motif, we computed the odds ratio and the significance of enrichment in each cluster, comparing to non-dynamic bin pool using Fisher’s exact test. The non-dynamic bin pool were sampled with replacement to match the distribution of average signal strength from the dynamic bins. Following that, significant TF PWMs were grouped in subfamilies using the structural information from TFClass^79^ because they share similar if not identical binding motifs. The top significantly over-represented TFs and their associated subfamilies were reported.

### A TAD-constrained map of enhancer-promoter associations

The reproducible strong enhancer calls (state #5) were merged using the mergeBed utility from BEDTools v2.17.0. After that, those regions or sub-regions overlapping the intervals +/− 2.5 kb from the TSS of genes in Gencode were excluded from the merged regions using subtractBed from BEDTools v2.17.0. Regions smaller than 2 kb were enlarged to 2 kb from their central coordinate (to allow for more robust signal estimation). This resulted in 66,556 putative enhancers. H3K27ac signals at these regions were then quantified using uniquely aligned, de-duplicated reads. These measurements were carried out by the coverageBed utility from BEDTools v2.17.0, then normalized to RPKM according to the sequencing depth of each sample, and log2-transformed (zeros were replaced by the smallest detectable value larger than zero). The mRNA expression of protein-coding genes was tracked across the 66 samples. Small and non-coding RNAs were excluded from any subsequent step by considering only those genes with a Gencode biotype supporting protein-coding functionality. FPKM were log2-transformed (zeros were replaced by the smallest detectable value larger than zero). For each TAD defined in the genome of mouse ES-cells^53^, the putative enhancers and genes were retrieved. All the enhancer-gene pairs within the TAD were then evaluated in terms of Spearman’s rank correlation coefficient (SCC) between the H3K27ac pattern of enrichment and the mRNA expression across the samples. Each gene was assigned to the putative enhancer showing the highest value of SCC. In order to attach p-values to these correlations, a null distribution was estimated empirically, by calculating the SCC of the enhancer with all the genes on the chromosome. Two different strategies were employed to estimate a p-value: 1) a z-score was calculated by subtracting the mean and dividing by the standard deviation of the null. The corresponding p-value was then calculated using the pnorm function in R; 2) an empirical p-value was defined as the number of times an equal or better than the observed SCC was found in the null. Only those putative enhancers showing a p-value <= 0.05 (for both strategies) and a SCC >= 0.25 were retained. Two maps were independently derived from the two biological replicates. These respectively encompassed 31,965 and 32,735 associations, with an overlap 21,142. Only these overlapping associations were used for further evaluation and analyses. **Validation of the enhancer-gene map using published chromatin conformation data**. Capture-C interaction data from the developing limb and brain^54^ were retrieved from the GEO (GSE84792). ChIA-PET interactions at sites bound by the cohesion subunit Smc1a in the developing limb^55^ were retrieved from the Supp. Table 2 of the original publication. Enhancer-gene contacts in fetal liver cells as inferred from Capture HiC^56^ were downloaded from ArrayExpress (E-MTAB-2414). In all cases, mm9 coordinates were mapped to mm10 using liftOver. For each published dataset, only those regions in the enhancer-gene map overlapping any experimentally validated interaction were retained. The fraction of interactions showing experimental support was then calculated for both the gene assigned by correlation and the nearest RefSeq gene. **Mapping of murine enhancer-gene map to human**. The putative enhancer regions were mapped to the human genome (hg19) using liftOver, with a strategy similar to previous reports^67^. Each region was required to both uniquely map to hg19, and to uniquely map back to the original region in mm10, with the requirement that >=50% of the bases in each region were mapped back to mouse after being mapped to human. For each enhancer-gene pair, the orthologous human gene was inferred using BioMart^82^. The orthologous pairs were also required to share the same TAD in human (TADs derived from human ES-cells^53^). **Validation of the enhancer-gene map using published eQTL-gene associations**. Single-tissue eQTL-gene associations generated by the GTEx consortium^83^ were downloaded from the GTEx portal (http://gtexportal.org, release v6p). Only those tissues with more than 750k annotated eQTLs were considered. A control set of enhancer-gene associations matching the size and the TSS-distance distributions of the real enhancer-gene map was generated. Briefly, for each enhancer-gene pair, the distance between the TSS of the gene and the central coordinate of the enhancer was calculated; after that, a region of the same size of the enhancer centered at the same distance to the TSS of the gene but on the opposite side of the enhancer was picked as a control set.

### Transgenic reporter assays

Names for functionally validated enhancers used throughout this work (mm numbers) are the unique identifiers from the VISTA Enhancer Browser (www.enhancer.lbl.gov)^65^. Enhancers were selected for testing as follows: The H3K27ac peak calls for three tissues (E12.5 heart, forebrain, and limb) were taken from the TSS-distal H3K27ac peaks called using the uniform processing pipeline (mm10-minimal) by the ENCODE Data Coordinating Center (narrow peaks from combined replicates). Peaks for each tissue were ranked by enrichment score (most to least significant). We then selected predicted enhancers from three different bins within each tissue’s ranked list for testing (bins were approximately ranks 1-85, 1500-1550, and 3000-3050). Loci that were already included in the VISTA Enhancer Browser or that appeared to overlap unannotated promoters were excluded from testing. In total, 150 predicted enhancers were tested, including 60 top ranked candidates (20 per tissue), 45 middle ranked (15 per tissue), and 45 lower ranked candidates (15 per tissue). Transgenic mouse assays were performed in FVB/NCrl strain *Mus musculus* animals (Charles River) as previously described^68,84^. Briefly, predicted enhancers were PCR amplified and cloned into a plasmid upstream of a minimal Hsp68 promoter and a *lacZ* reporter gene. Transgenic embryos were generated by pronuclear injection of the resulting plasmids into fertilized mouse eggs. Embryos were implanted into surrogate mothers, collected at E12.5, and stained for β-galactosidase activity. Elements were scored as positive enhancers if at least three embryos had identical β-galactosidase staining in the same tissue. Elements were scored as negative if no reproducible staining was observed and at least five embryos harboring a transgene insertion were obtained. Genomic coordinates and results for each element are provided in Supplementary Table 5, through the ENCODE project data portal (www.encodeproject.org), and at the VISTA Enhancer Browser website (www.enhancer.lbl.gov).

### Mapping to repeat element families

Since ENCODE analysis pipeline were focusing on the uniquely mapped reads, a lot of information provided by the reads that are not uniquely mapped will be missed. This is especially undesirable for repetitive element analysis as there is a higher chance for read coming from a repetitive region of DNA will map un-uniquely to the reference genome. Hence, we use a new pipeline with two rounds of mapping steps to re-process all the fastq files. The pipeline utilizes all the reads for the purpose of gaining enough sensitivity to uncover the enrichment of any repetitive element subfamilies that were previously undiscovered because of low coverage. In the first round of mapping, sample reads were aligned to the reference genome mm10 using Bowtie1 with: bowtie hg19 -p 16 -t -m 1 -S –chunkmbs 512 –max multimap.fastq input.fastq output.sam^85^. –max is used to separate reads mapping to multiple locations of the genome from uniquely mapped reads. In the second round of mapping, a customized assemblies file are constructed by concatenating genomic instances of each repetitive element subfamily, their 15 bp flanking genomic sequences and a 200bp spacer sequence in FASTA format^86^. The annotation file for repetitive elements used in this step was downloaded from Repeatmasker.org. A python script used with parameters are as following: python RepEnrich.py/data/mm10_repeatmasker.txt/data/sample_A sample_A/data/mm10_setup_folder sampleA_multimap.fastq sampleA_unique.bam –cpus 16^87^. The number of reads mapping to repetitive element subfamilies, repetitive element families, or repetitive element classes are determined using information from both uniquely mapped reads that overlap with repetitive element and un-uniquely mapped reads. Since some of the repetitive element subfamilies are very similar to each other, a fractional counts method was used to classify the reads that map to multiple repetitive element subfamilies. It sums reads mapping uniquely to a repetitive element subfamily once and counts reads mapping to multiple subfamilies using a fraction 1/Ns, where Ns = number of repetitive element subfamilies the read aligns with. A table of counts that estimate enrichment signal for the repeats classes across different tissues is built as the final output for plotting the figures.

### Data processing in R

Most of the described data processing steps (plotting, statistical tests, calculating correlations and hierarchical clustering) were performed in the statistical computing environment R v.3.3.1 (www.r-project.org).

## DATA ACCESS

Raw and processed ChIP-seq data reported here can be accessed via the ENCODE Data Collection and Coordiantion (DCC) website www.encodedcc.org. More specifically, a full list of the ChIP-seq experiments can be found here: https://www.encodeproject.org/search/?type=Experiment&assay_slims=DNA+binding&assay_title=ChIP-seq&award.rfa=ENCODE3&lab.title=Bing+Ren%2C+UCSD&limit=all. ChromHMM data can be found here: http://enhancer.sdsc.edu/enhancer_export/ENCODE/

## EXTENDED DATA FIGURE LEGENDS

**Extended data figure 1. Summary of primary ChIP-seq data.** (a) Table summarizing the characteristic enrichment patterns for each histone modification surveyed here. Note that modifications are generally categorized as “narrow” or “broad” depending on the typical breadth of enrichment. H3K9me3 is further distinguished from other broad marks because it shows very few regions of enrichment in non-repetitive sequence in primary tissues and cells^42^. (b) Metagene profiles illustrating the typical patterns of histone modification enrichment at active genes (defined as RPKM > 10 in all tissue-stages surveyed here). ChIP-seq data plotted is from embryonic Heart at E13.5. (c) Sequencing depth is plotted for every library reported (N=1,128 ChIP + 168 input). ENCODE “usable” read depth standards (mapq>30, and after PCR duplicate removal) are indicated to the right. Note that read depth standards changed part way through our study: increasing from 10M to 20M for narrow marks, 20M to 45M for broad marks, and 10M to 30M for input. All narrow mark libraries exceed the 10M minimal depth. Broad mark libraries exceed the 20M minimal depth with only 4 exceptions, all of which exceed 19M. Input libraries exceed the 10M minimal depth with only 1 exception, which exceed 9.7M. The read depth standard for H3K9me3 >45M mapped reads of any mapq (because most H3k9me3 is repetitive sequence, see Extended Figure 6); all H3K9me3 libraries exceed this threshold. (d) Mapping quality is plotted for every library, measured as the fraction of reads with mapping quality scores (mapq) > 30. Reads with lower mapq scores (i.e. non-uniquely mapping reads) were eliminated from downstream analysis. (e) Three metrics of library complexity are plotted (NRF, PBC1, PBC2). See ENCODE data standards^88^ for detailed descriptions and formulas. Tables below each plot show the percent of libraries that exceed the thresholds indicated. (f) Two measures of signal-to-noise ration are plotted (NSC, RSC). Again, detailed descriptions are available in the ENCODE data standards descriptions.

**Extended data figure 2. ChIP-seq peak calling.** (a) Schematic representation of ChIP-seq peak calling pipeline. More information can be found here: https://www.encodeproject.org/chip-seq/. (b) Four different peak summary statistics are plotted for every tissue-stage. From top to bottom: 1) Number of peaks called (passing IDR threshold); 2) Total coverage of those peaks; 3) Peak coverage as in (2), but separated according to tissue; 4) Peak coverage as in (2), but separated by stage. Note that E10.5 ChIP-seq experiments were performed with a modified protocol, and in some cases a different more sensitive antibody (H3K27ac, H3K4me1). We suspect that is why E10.5 sometimes appears as an outlier in terms of coverage. (c) Peak reproducibility as measured by the percentage of peaks called form the pooled data that were called independently in both individual replicates. (d) Peak reproducibility as measured by correlation of peak strengths (average bigwig signal within each peak) between independent replicates. (e) Principle Component Analysis of all tissue-stages based on the distribution of ChIP-seq signal across the genome.

**Extended data figure 3. Dynamic ChIP-seq signal across developmental space and time.** (a) Genome browser view of *Gad1* (chr2:70,547,104-70,615,401; mm10), a marker of GABAergic neurons, showing the gain of active chromatin signatures during forebrain development. (b-c) Genome browser views of *Ccnb1* (chr13:100,776,802-100,788,423; mm10) and Cdk2 (chr10:128,693,493-128,709,497; mm10), key cell cycle regulators, showing the loss of active chromatin signatures during forebrain development. (d) multi-tiered plot showing the relationship between tissue-specificity, stage-specificity, and location relative to TSS. For each mark, peaks were merged across all tissue-stages. These merged peaks were then overlapped with each tissue-stage individually to determine how many individual tissues and stages that peak was called in. The x-axis at the top indicates how many tissues a given peak was called in (1 - 12). The top line graph shows the tissue-specificity of peaks, as the percentage of total peaks for a given mark that were called in 1 - 12 tissues. The middle heatmap shows stage specificity, as the fraction of stages within a given tissue that a peak is called in. In other words, if a peak is called in 3 tissues, what fraction of the available stages within those 3 tissues was the peak called in. Note that peaks which are more restricted to specific tissues, are also more restricted to specific stages within those tissues. The bottom heatmap shows the location of peaks relative to TSS, as the fraction of peaks that overlap an annotated GENCODE TSS. Note that peaks which are more consistent across tissues and across stages are much more likely to overlap a TSS. (e) Heapmap showing the H3K27ac ChIP-seq signal at H3K27ac ChIP-seq peaks called in forebrain (merged across stages). Peaks are clustered based on how many stages within forebrain they were called in (y axis, left). The number of peaks in each cluster is indicated to the right.

**Extended data figure 4. 15-state chromHMM model.** (a) Schematic representation of the chromatin states identification strategy applied in this study. (b) Heat maps showing the maximum correlation of each state in the “full” model (y-axis) with its best matching state in each simpler model (x-axis). The median correlation of all the 24 state is shown in the plots on top of the heat maps. (c) Plot showing the classification of the k-means clustering of the emission probabilities from all the models. The optimal number of states was defined by the smallest value of k showing a ratio equal or higher than 95% (orange line) of the maximum clusters’ separation (red line). SS = Sum of Squares. (d) Heat maps showing the emission probabilities for each chromatin mark in each state, as defined by ChromHMM, for both replicates. (e) Box plots showing the similarity between replicates, among samples from the same tissue and among all samples, considering either active TSS (state #1, red) or strong enhancers (state #5, orange). P-values from Mann-Whitney tests. Similarity measured as pairwise binary distance (see Methods). (f) Bar plot showing the enrichment of each mark in state 11 (permissive) relative to state 15 (no signal, genomic background). Note that the chromHMM emission probability for H3K36me3 in state 11 is >30-fold higher than genomic background. (g) Comparison of the ChromHMM model reported here with previously published chromHMM models. Grey vertical bars indicate histone modifications profiled here that were not included in those previous studies. Horizontal bars indicate chromatin states identified in our study that do not have a clear counterpart in those studies.

**Extended data figure 5. Chromatin states at developmental genes.** (a) Genome browser views showing tissue-restricted activity patterns at *Nkx2-5* (chr17:26,818,483-26,870,007), *Cdx2* (chr5:147,294,550-147,313,599), *Barx1* (chr13:48,649,148-48,680,395), *Wt1* (chr2:105,097,427-105,200,306), and *Foxg1* (chr12:49,362,757-49,416,342). *Myc* as shown in the far right as an example of a gene that is not tissue restricted across our developmental series (chr15:61,972,107-62,003,584). All coordinates are mm10. (b) Genome browser views showing stage-restricted activity patterns at *NeuroD2* (chr11:98,308,801-98,345,075) and *Gad1* (chr2:70,547,104-70,615,401). (c) This plot is a counterpart to Figure 4b. The relative frequency of active chromatin (State 1) at four different sets of TSS: 1) all GENCODE protein coding TSS (N=90,384), 2) the subset of (1) that belong to transcripts of TF genes (N=1,019), 3) the subset of (2) that are also known to cause to Mendelian phenotypes when mutated (i.e. “OMIM” genes; N=227), 4) the remaining OMIM gene TSS that do not encode TFs (N=3,497). The individual gene symbols indicate genes that are known to cause congenital defects when abnormally expressed. Not shown: TSS active in none of the 66 tissue-stages (43%, 32 %, 33%, and 39%, respectively), and TSS active in all 66 tissue stages (14%, 19%, 15%, and 13 %, respectively).

**Extended data figure 6. H3K9me3 marked heterochromatin.** (a) Genome browser view showing a large region of chromosome 15 (chr15:87,165,311-104,043,685; mm10). Signal tracks (fold enrichment over input) are shown for all marks. Note that H3K9me3 looks relatively flat, unlike the other marks. Note also that this signal does not include reads that map non-uniquely to the genome (i.e. map to repetitive elements, mapq<30). We find very few regions of strong H3K9me3 enrichment outside of repetitive elemetns, consistent with previous reports of H3K9me3 distribution in primary tissues^42^. (b) Bar chart showing the fraction of total sequencing reads that mapped to the reference genome (light green), and that mapped uniquely to the reference genome (mapq >= 30; dark green). Bar height represents the mean for all ChIP- libraries reported here separated by mark, and error bars represent standard deviation. Control bars represent ChIP input libraries (no IP step). Note that all marks and input have a high mapping rate (mean >90%). However, H3K9me3 has a markedly low rate of unique mapping, suggesting that this modification is specifically enriched in non-unique (i.e. repetitive) genomic regions. (c) Stacked bar plots show the type of repetitive elements that the non-uniquely mapping reads from (b) correspond to (see methods). Note that H3K9me3 reads are highly enriched in satellite repeats relative to the input control samples. (d) Genome browser view of chromatin states showing that some *Pchd* (chr18:36,720,767-38,058,585; mm10) and *Zfp* (chr11:50,774,724-50,939,391; mm10) genes do show significant H3K9me3 enrichment (state 14) during development. 3’ UTRs of Zfp genes marked by H3K9me3 (noted previously^44^) are indicated by pink arrow heads. (e) Similar to (d), but showing raw ChIP-seq fold enrichment tracks for each mark across these regions.

**Extended data figure 7. Dynamic chromatin states.** (a) Heatmap shows the fraction of bases for each state in forebrain that are in a different state in other tissues at the same stage. The bar plot on the right shows the median value (fraction of variable bases) considering all tissues. Enhancer states, which are highlighted by a vertical bar, are the most variable between tissues. (b) Dendrogram showing the hierarchical relationships among strong enhancers (state #5) in different tissues during development (clustering according to binary distance, Ward’s method). (c) Same as (b) but considering only limb and face samples. (d) Heat map showing the statistical significance of the enriched functional terms (GO Biological Process, as assessed by GREAT, x axis) for the sets of strong enhancers (state #5) across each tissue-stage (y axis). The terms were hierarchically clustered (average linkage) according to Pearson’s correlation. A subset of the terms highly enriched in both Limb and Face are listed below the main heatmap. (e) Heatmap shows the fraction of bases for each state in forebrain e11.5 that are in a different state at other time points. The bar plot on the right shows the median value (fraction of variable bases) considering all tissues. (f) Heat map showing the most enriched biological processes (GO terms) for the genes in the vicinity of liver enhancers. For each one of the indicated sets, the significantly enriched terms were identified and divided into deciles (based on statistical significance). The ten most enriched terms for each set were then grouped together and hierarchically clustered. Genes involved in either hematopoiesis or metabolic processes are color-coded as indicated. (g) Sankey diagram showing the tendencies of each chromatin state at a given stage (left, Stage N) to transition to other chromatin states at the following developmental stage (right, Stage N+1).

**Extended data figure 8. Dynamic enhancer patterns for 12 tissues.** K-means clustering of dynamic enhancers in each tissue based on H3K27ac signal at available stage from E11.5 to P0. The number of clusters within each tissue, and the number of dynamic enhancers within each cluster, are indicated to the left of each heat map. Corresponding GO enrichment and motifs are provided in Table S1.

**Extended data figure 9. Predicting enhancer target genes.** (a) Schematic of the approach to assign enhancers to target genes. Correlation between H3K27ac signal at putative enhancers (green bars) and mRNA expression of genes (black bars) in the same TAD across tissues and developmental time was used to infer the most likely enhancer-gene interactions. (b) Genome browser view showing chromatin states at the Ascl1 locus, as in Figure 5a, but showing ChIP-seq fold enrichment tracks instead of chromatin states. (c) (Upper panel) Plot showing the fraction of enhancer-gene associations common to the two replicates (Reproducibility) as a function of the stringency applied to the Spearman’s Correlation Coefficient (SCC). The two curves correspond to developmental time series. (Lower panel) Scatterplots showing the relation between the number of putative enhancers per gene at the default cutoff (SCC = 0.25, x-axis) vs a more stringent cutoff (SCC = 0.45 or 0.75, respectively, y-axis). (d) Scatterplot showing a comparison of the number of putative enhancers per gene in the two replicates. (e) Histogram showing the distribution of the number of enhancers per gene. (f) The most significantly enriched GO terms are listed for genes that have one predicted enhancer (top) and genes with >= 5 predicted enhancer (bottom). (g) As in Figure 5b, this plot shows that enhancer-gene interactions identified by this correlative approach are generally more likely to be supported by chromatin interaction data than associations derived by a nearest gene approach. To ensure that this was not due to an artifact of the chromatin capture technologies being unable to detect very short range interactions, we used different distance cutoffs (10Kb, 100Kb) to define the “nearest” non-target gene. (h) Bar plot show the frequency of GWAS hits (EMBL-EBI catalog) in three different types of enhancers: enhancers predicted to target OMIM genes (top); enhancers predicted to target genes other than OMIM genes (middle); enhancers in the same TADs as OMIM genes, but not predicted to target those OMIM genes (bottom).

**Extended data figure 10. Transgenic validation results for predicted enhancers.** (a) Heat map showing the tissue-specific enrichments of VISTA enhancers for the different chromatin states in E11.5 heart, limb and forebrain. (b) Representative e12.5 transgenic embryos for each of the 61 enhancers that validated in the expected tissue (forebrain, limb, or heart). Reporter gene expression is indicated by blue staining, and enhancer names (mm numbers) are the unique identifiers from the VISTA Enhancer Browser^65^. Red arrows indicate the position of the forebrain, limbs, or heart. See also Supplementary Table 5 for results. (c) Violin plots showing transgenic enhancer assay reproducibility for different rank classes of tested elements. Only those enhancers that validated in the correct expected tissue are shown. Reproducibility differences between rank classes were not statistically significant (Mann-Whitney U Test).

## REFERENCES

1 Allis, C. D. & Jenuwein, T. The molecular hallmarks of epigenetic control. Nature Reviews Genetics 17, 487-500, doi:doi:10.1038/nrg.2016.59 (2016).

2 Bannister, A. J. & Kouzarides, T. Regulation of chromatin by histone modifications. Cell Research 21, 381-395, doi:doi:10.1038/cr.2011.22 (2011).

3 Tessarz, P. & Kouzarides, T. Histone core modifications regulating nucleosome structure and dynamics. Nature reviews. Molecular cell biology 15, 703-708, doi:10.1038/nrm3890 (2014).

4 Campos, E. I., Stafford, J. M. & Reinberg, D. Epigenetic inheritance: histone bookmarks across generations. Trends in cell biology 24, 664-674, doi:10.1016/j.tcb.2014.08.004 (2014).

5 Fahrner, J. A. & Bjornsson, H. T. Mendelian disorders of the epigenetic machinery: tipping the balance of chromatin states. Annual review of genomics and human genetics 15, 269-293, doi:10.1146/annurev-genom-090613-094245 (2014).

6 Krumm, N., O’Roak, B. J., Shendure, J. & Eichler, E. E. A de novo convergence of autism genetics and molecular neuroscience. Trends in neurosciences 37, 95-105, doi:10.1016/j.tins.2013.11.005 (2014).

7 Roy, D. M., Walsh, L. A. & Chan, T. A. Driver mutations of cancer epigenomes. Protein & cell 5, 265-296, doi:10.1007/s13238-014-0031-6 (2014).

8 Bernstein, B. E., Meissner, A. & Lander, E. S. The mammalian epigenome. Cell 128, 669-681, doi:10.1016/j.cell.2007.01.033 (2007).

9 Rivera, C. M. & Ren, B. Mapping human epigenomes. Cell 155, 39-55, doi:10.1016/j.cell.2013.09.011 (2013).

10 Heintzman, N. D. et al. Distinct and predictive chromatin signatures of transcriptional promoters and enhancers in the human genome. Nature genetics 39, 311-318, doi:10.1038/ng1966 (2007).

11 Rada-Iglesias, A. et al. A unique chromatin signature uncovers early developmental enhancers in humans. Nature 470, 279-283, doi:10.1038/nature09692 (2011).

12 Creyghton, M. P. et al. Histone H3K27ac separates active from poised enhancers and predicts developmental state. Proceedings of the National Academy of Sciences of the United States of America 107, 21931-21936, doi:10.1073/pnas.1016071107 (2010).

13 Ernst, J. & Kellis, M. in Nature methods Vol. 9 215-216 (2012).

14 Ernst, J. et al. Mapping and analysis of chromatin state dynamics in nine human cell types. Nature 473, 43-49, doi:10.1038/nature09906 (2011).

15 Hon, G., Ren, B. & Wang, W. ChromaSig: a probabilistic approach to finding common chromatin signatures in the human genome. PLoS computational biology 4, e1000201, doi:10.1371/journal.pcbi.1000201 (2008).

16 Kundaje, A. et al. Integrative analysis of 111 reference human epigenomes. Nature 518, 317-330, doi:10.1038/nature14248 (2015).

17 GmbH, E. Reference Epigenome Standards · IHEC, <http://ihecepigenomes.org/research/reference-epigenome-standards/> (2017).

18 Li, Q., Brown, J. B., Huang, H. & Bickel, P. J. Measuring reproducibility of high-throughput experiments. doi:10.1214/11-AOAS466 (2011).

19 England, J. & Loughna, S. Heavy and light roles: myosin in the morphogenesis of the heart. Cellular and molecular life sciences: CMLS 70, 1221-1239, doi:10.1007/s00018-012-1131-1 (2013).

20 McCormick, M. B. et al. NeuroD2 and neuroD3: distinct expression patterns and transcriptional activation potentials within the neuroD gene family. Molecular and cellular biology 16, 5792-5800 (1996).

21 Kodama, T. et al. Neuronal classification and marker gene identification via single-cell expression profiling of brainstem vestibular neurons subserving cerebellar learning. J Neurosci 32, 7819-7831, doi:10.1523/jneurosci.0543-12.2012 (2012).

22 Noonan, J. P. & McCallion, A. S. Genomics of long-range regulatory elements. Annual review of genomics and human genetics 11, 1-23, doi:10.1146/annurev-genom-082509-141651 (2010).

23 Visel, A., Rubin, E. M. & Pennacchio, L. A. Genomic views of distant-acting enhancers. Nature 461, 199-205, doi:10.1038/nature08451 (2009).

24 Yue, F. et al. A comparative encyclopedia of DNA elements in the mouse genome. Nature 515, 355-364, doi:10.1038/nature13992 (2014).

25 Barski, A. et al. High-resolution profiling of histone methylations in the human genome. Cell 129, 823-837, doi:10.1016/j.cell.2007.05.009 (2007).

26 McCulley, D. J. & Black, B. L. Transcription factor pathways and congenital heart disease. Current topics in developmental biology 100, 253-277, doi:10.1016/b978-0-12-387786-4.00008-7 (2012).

27 Costa, R. H., Kalinichenko, V. V. & Lim, L. Transcription factors in mouse lung development and function. American journal of physiology. Lung cellular and molecular physiology 280, L823-838 (2001).

28 Sheaffer, K. L. & Kaestner, K. H. Transcriptional Networks in Liver and Intestinal Development. doi:10.1101/cshperspect.a008284 (2012).

29 Jayewickreme, C. D. & Shivdasani, R. A. Control of stomach smooth muscle development and intestinal rotation by transcription factor BARX1. Developmental biology 405, 21-32, doi:10.1016/j.ydbio.2015.05.024 (2015).

30 Dressler, G. R. Transcription factors in renal development: the WT1 and Pax-2 story. Seminars in nephrology 15, 263-271 (1995).

31 Bantignies, F. & Cavalli, G. Polycomb group proteins: repression in 3D. Trends in genetics: TIG 27, 454-464, doi:10.1016/j.tig.2011.06.008 (2011).

32 Piunti, A. & Shilatifard, A. Epigenetic balance of gene expression by Polycomb and COMPASS families. Science 352, aad9780, doi:10.1126/science.aad9780 (2016).

33 Aloia, L., Stefano, B. D. & Croce, L. D. Polycomb complexes in stem cells and embryonic development. doi:10.1242/dev.091553 (2013).

34 McLean, C. Y. et al. GREAT improves functional interpretation of cis-regulatory regions. Nature biotechnology 28, 495-501, doi:10.1038/nbt.1630 (2010).

35 Online Mendelian Inheritance in Man, O. McKusick-Nathans Institute of Genetic Medicine, Johns Hopkins University (Baltimore, MD). (2017).

36 Franke, M. et al. Formation of new chromatin domains determines pathogenicity of genomic duplications. Nature 538, 265-269, doi:10.1038/nature19800 (2016).

37 Lupianez, D. G. et al. Disruptions of topological chromatin domains cause pathogenic rewiring of gene-enhancer interactions. Cell 161, 1012-1025, doi:10.1016/j.cell.2015.04.004 (2015).

38 Lettice, L. A. et al. Disruption of a long-range cis-acting regulator for Shh causes preaxial polydactyly. Proceedings of the National Academy of Sciences of the United States of America 99, 7548-7553, doi:10.1073/pnas.112212199 (2002).

39 Spitz, F. & Furlong, E. E. Transcription factors: from enhancer binding to developmental control. Nature reviews. Genetics 13, 613-626, doi:10.1038/nrg3207 (2012).

40 Saksouk, N., Simboeck, E. & Dejardin, J. Constitutive heterochromatin formation and transcription in mammals. Epigenetics & chromatin 8, 3, doi:10.1186/1756-8935-8-3 (2015).

41 Bulut-Karslioglu, A. et al. Suv39h-dependent H3K9me3 marks intact retrotransposons and silences LINE elements in mouse embryonic stem cells. Molecular cell 55, 277-290, doi:10.1016/j.molcel.2014.05.029 (2014).

42 Zhu, J. et al. Genome-wide chromatin state transitions associated with developmental and environmental cues. Cell 152, 642-654, doi:10.1016/j.cell.2012.12.033 (2013).

43 Lehnertz, B. et al. Suv39h-mediated histone H3 lysine 9 methylation directs DNA methylation to major satellite repeats at pericentric heterochromatin. Current biology: CB 13, 1192-1200 (2003).

44 Blahnik, K. R. et al. Characterization of the contradictory chromatin signatures at the 3′ exons of zinc finger genes. PloS one 6, e17121, doi:10.1371/journal.pone.0017121 (2011).

45 Kaucka, M. et al. Analysis of neural crest-derived clones reveals novel aspects of facial development. Science advances 2, e1600060, doi:10.1126/sciadv.1600060 (2016).

46 Gillis, J. A. & Hall, B. K. A shared role for sonic hedgehog signalling in patterning chondrichthyan gill arch appendages and tetrapod limbs. Development 143, 1313-1317, doi:10.1242/dev.133884 (2016).

47 Gordillo, M., Evans, T. & Gouon-Evans, V. Orchestrating liver development. doi:10.1242/dev.114215 (2015).

48 Nord, A. S. et al. Rapid and pervasive changes in genome-wide enhancer usage during mammalian development. Cell 155, 1521-1531, doi:10.1016/j.cell.2013.11.033 (2013).

49 Avilion, A. A. et al. Multipotent cell lineages in early mouse development depend on SOX2 function. Genes & development 17, 126-140, doi:10.1101/gad.224503 (2003).

50 Shimozaki, K. Sox2 transcription network acts as a molecular switch to regulate properties of neural stem cells. World J Stem Cells 6, 485-490, doi:10.4252/wjsc.v6.i4.485 (2014).

51 Uittenbogaard, M., Baxter, K. K. & Chiaramello, A. NeuroD6 genomic signature bridging neuronal differentiation to survival via the molecular chaperone network. Journal of neuroscience research 88, 33-54, doi:10.1002/jnr.22182 (2010).

52 Barbosa, A. C. et al. MEF2C, a transcription factor that facilitates learning and memory by negative regulation of synapse numbers and function. doi:10.1073/pnas.0802679105 (2008).

53 Dixon, J. R. et al. Topological domains in mammalian genomes identified by analysis of chromatin interactions. Nature 485, 376-380, doi:10.1038/nature11082 (2012).

54 Andrey, G. et al. Characterization of hundreds of regulatory landscapes in developing limbs reveals two regimes of chromatin folding. Genome research 27, 223-233, doi:10.1101/gr.213066.116 (2017).

55 DeMare, L. E. et al. The genomic landscape of cohesin-associated chromatin interactions. Genome research 23, 1224-1234, doi:10.1101/gr.156570.113 (2013).

56 Schoenfelder, S. et al. The pluripotent regulatory circuitry connecting promoters to their long-range interacting elements. Genome research 25, 582-597, doi:10.1101/gr.185272.114 (2015).

57 The Genotype-Tissue Expression (GTEx) project. Nature genetics 45, 580-585, doi:10.1038/ng.2653 (2013).

58 Gaffney, D. J. et al. Dissecting the regulatory architecture of gene expression QTLs. Genome biology 13, R7, doi:10.1186/gb-2012-13-1-r7 (2012).

59 Sobreira, N. L. & Valle, D. Lessons learned from the search for genes responsible for rare Mendelian disorders. Molecular genetics & genomic medicine 4, 371-375, doi:10.1002/mgg3.233 (2016).

60 Maurano, M. T. et al. Systematic localization of common disease-associated variation in regulatory DNA. Science 337, 1190-1195, doi:10.1126/science.1222794 (2012).

61 Kathiresan, S. & Srivastava, D. Genetics of Human Cardiovascular Disease. Cell 148, 1242-1257, doi:10.1016/j.cell.2012.03.001 (2012).

62 Smemo, S. et al. Obesity-associated variants within FTO form long-range functional connections with IRX3. Nature 507, 371-375, doi:10.1038/nature13138 (2014).

63 Canver, M. C. et al. BCL11A enhancer dissection by Cas9-mediated in situ saturating mutagenesis. Nature 527, 192-197, doi:doi:10.1038/nature15521 (2015).

64 Hnisz, D. et al. Super-enhancers in the control of cell identity and disease. Cell 155, 934-947, doi:10.1016/j.cell.2013.09.053 (2013).

65 Visel, A., Minovitsky, S., Dubchak, I. & Pennacchio, L. A. VISTA Enhancer Browser–a database of tissue-specific human enhancers. Nucleic acids research 35, D88-92, doi:10.1093/nar/gkl822 (2007).

66 Shen, Y. et al. A map of the cis-regulatory sequences in the mouse genome. Nature 488, 116-120, doi:10.1038/nature11243 (2012).

67 Cotney, J. et al. Chromatin state signatures associated with tissue-specific gene expression and enhancer activity in the embryonic limb. Genome research 22, 1069-1080, doi:10.1101/gr.129817.111 (2012).

68 Kothary, R. et al. Inducible expression of an hsp68-lacZ hybrid gene in transgenic mice. Development 105, 707-714 (1989).

69 Thurman, R. E. et al. The accessible chromatin landscape of the human genome. Nature 489, 75-82, doi:10.1038/nature11232 (2012).

70 Dickel, D. E. et al. Genome-wide compendium and functional assessment of in vivo heart enhancers. Nature communications 7, 12923, doi:10.1038/ncomms12923 (2016).

71 Li, Y. et al. CRISPR reveals a distal super-enhancer required for Sox2 expression in mouse embryonic stem cells. PloS one 9, e114485, doi:10.1371/journal.pone.0114485 (2014).

72 Attanasio, C. et al. Fine tuning of craniofacial morphology by distant-acting enhancers. Science 342, 1241006, doi:10.1126/science.1241006 (2013).

73 Sloan, C. A. et al. ENCODE data at the ENCODE portal. Nucleic acids research 44, D726-732, doi:10.1093/nar/gkv1160 (2016).

74 Marinov, G. K., Kundaje, A., Park, P. J. & Wold, B. J. Large-scale quality analysis of published ChIP-seq data. G3 (Bethesda, Md.) 4, 209-223, doi:10.1534/g3.113.008680 (2014).

75 Ramirez, F., Dundar, F., Diehl, S., Gruning, B. A. & Manke, T. deepTools: a flexible platform for exploring deep-sequencing data. Nucleic acids research 42, W187-191, doi:10.1093/nar/gku365 (2014).

76 Quinlan, A. R. & Hall, I. M. BEDTools: a flexible suite of utilities for comparing genomic features. Bioinformatics 26, 841-842, doi:10.1093/bioinformatics/btq033 (2010).

77 Wang, J., Duncan, D., Shi, Z. & Zhang, B. WEB-based GEne SeT AnaLysis Toolkit (WebGestalt): update 2013. Nucleic acids research 41, W77-83, doi:10.1093/nar/gkt439 (2013).

78 Harrow, J. et al. GENCODE: the reference human genome annotation for The ENCODE Project. Genome research 22, 1760-1774, doi:10.1101/gr.135350.111 (2012).

79 Wingender, E., Schoeps, T. & Donitz, J. TFClass: an expandable hierarchical classification of human transcription factors. Nucleic acids research 41, D165-170, doi:10.1093/nar/gks1123 (2013).

80 Ritchie, M. E. et al. limma powers differential expression analyses for RNA-sequencing and microarray studies. Nucleic acids research 43, e47, doi:10.1093/nar/gkv007 (2015).

81 Bailey, T. L. et al. MEME SUITE: tools for motif discovery and searching. Nucleic acids research 37, W202-208, doi:10.1093/nar/gkp335 (2009).

82 Smedley, D. et al. The BioMart community portal: an innovative alternative to large, centralized data repositories. Nucleic acids research 43, W589-598, doi:10.1093/nar/gkv350 (2015).

83 Human genomics. The Genotype-Tissue Expression (GTEx) pilot analysis: multitissue gene regulation in humans. Science 348, 648-660, doi:10.1126/science.1262110 (2015).

84 Pennacchio, L. A. et al. In vivo enhancer analysis of human conserved non-coding sequences. Nature 444, 499-502, doi:10.1038/nature05295 (2006).

85 Langmead, B., Trapnell, C., Pop, M. & Salzberg, S. L. Ultrafast and memory-efficient alignment of short DNA sequences to the human genome. Genome biology 10, R25, doi:10.1186/gb-2009-10-3-r25 (2009).

86 Day, D. S., Luquette, L. J., Park, P. J. & Kharchenko, P. V. Estimating enrichment of repetitive elements from high-throughput sequence data. Genome biology 11, R69, doi:10.1186/gb-2010-11-6-r69 (2010).

87 Criscione, S. W., Zhang, Y., Thompson, W., Sedivy, J. M. & Neretti, N. Transcriptional landscape of repetitive elements in normal and cancer human cells. BMC genomics 15, 583, doi:10.1186/1471-2164-15-583 (2014).

88 Landt, S. G. et al. ChIP-seq guidelines and practices of the ENCODE and modENCODE consortia. Genome research 22, 1813-1831, doi:10.1101/gr.136184.111 (2012).

